# The microenvironment dictates glycocalyx construction and immune surveillance

**DOI:** 10.1101/2023.06.23.546317

**Authors:** Kevin M. Tharp, Sangwoo Park, Greg A. Timblin, Alicia L. Richards, Jordan A. Berg, Nicholas M. Twells, Nicholas M. Riley, Egan L. Peltan, D. Judy Shon, Erica Stevenson, Kimberly Tsui, Francesco Palomba, Austin E. Y. T. Lefebvre, Ross W. Soens, Nadia M.E. Ayad, Johanna ten Hoeve-Scott, Kevin Healy, Michelle Digman, Andrew Dillin, Carolyn R. Bertozzi, Danielle L. Swaney, Lara K. Mahal, Jason R. Cantor, Matthew J. Paszek, Valerie M. Weaver

## Abstract

Efforts to identify anti-cancer therapeutics and understand tumor-immune interactions are built with *in vitro* models that do not match the microenvironmental characteristics of human tissues. Using *in vitro* models which mimic the physical properties of healthy or cancerous tissues and a physiologically relevant culture medium, we demonstrate that the chemical and physical properties of the microenvironment regulate the composition and topology of the glycocalyx. Remarkably, we find that cancer and age-related changes in the physical properties of the microenvironment are sufficient to adjust immune surveillance via the topology of the glycocalyx, a previously unknown phenomenon observable only with a physiologically relevant culture medium.

**Key Points:**

- Culture medium dictates cellular mechanoresponse signatures in vitro
- Epithelial glycocalyx construction is mediated by Heat Shock Factor 1 (HSF1)
- Sialic acid topology dictates Natural Killer cell cytotoxicity
- Physiological microenvironments reveal distinct glycobiology

## Introduction

The physical properties of the microenvironment determine cellular phenotypes and functions in healthy and diseased tissues (*1*–*3*). Cell-extrinsic mechanical properties affect adhesion and cytoskeletal dynamics, which have an emerging role in the regulation of metabolism (*4*–*9*). Mechanical regulation of metabolism can reciprocally tune the physical properties of cells and tissues by rewiring metabolism to synthesize structural macromolecules like the extracellular matrix (ECM) (*10*) or glycocalyx (*11*–*13*), a pericellular matrix composed of glycoproteins and glycolipids that is especially apparent on the surface of endothelial and epithelial cells. This type of “mechanoreciprocity” may serve to buffer mechanical stresses by reducing the need for energetically demanding cytoskeletal responses in favor of passive physical buffering via the ECM or glycocalyx (*14*, *15*). However, stress management strategies can have unintended consequences and adaptive changes to the glycocalyx or ECM can dysregulate immune surveillance (*16*–*19*).

## Results

Recent studies have demonstrated that cell-intrinsic or -extrinsic physical properties affect cellular metabolism (*4*, *5*, *9*, *20*–*23*). However, these studies were carried out in conventional *in vitro* culture media (e.g., Dulbecco’s Modified Eagle Medium, DMEM) that can obfuscate physiologically relevant metabolic wiring and regulation because they do not model *in vivo* nutrient availability (*24*–*27*). This led us to hypothesize that human cells may differentially respond to the physical properties of the microenvironment (“mechanoresponse”) in a culture medium systematically designed to more faithfully reflect the nutrient composition of human plasma (Human Plasma-Like Medium, HPLM) rather than a conventional culture medium (DMEM) (Fig. S1A-C). Typically, epithelial cells exposed to stiff adhesive substrates use actomyosin-supported cell-matrix adhesions to spread across the surface, whereas cells on soft substrates enforce cell-cell junctions and remain in colonies. Fluorescence microscopy revealed that these morphological dynamics were observable in both media (Fig. S1D), but cortical actin networks appeared more elaborate and ruffled in HPLM.

### Proteome dynamics and the microenvironment

To test how medium composition impacts cellular mechanoresponses, we used mass spectrometry-based proteomics to compare cells cultured on soft substrates that recapitulate the elastic modulus of the normal mammary stroma (400 Pa) to cells cultured on stiff substrates that recapitulate the elastic modulus of malignant tumors (60k Pa) (Fig. 1A). As expected, we found that ECM proteins such as fibronectin (FN1) were enriched in cells cultured on stiff substrates. However, the proteomic changes induced by substrate elasticity showed a limited correlation between the two media (Fig. 1B-D and S1E-F). To visualize this relationship, we overlaid proteomic changes affected by adhesion substrate stiffness observed in DMEM onto the data obtained in HPLM (red or blue boxes on the volcano plot) (Fig. 1E-F). This revealed that the lack of correlation between mechanoresponses in HPLM versus DMEM was attributed to differences in the directionality and magnitude of either enriched or reduced proteins.

**Figure 1:**
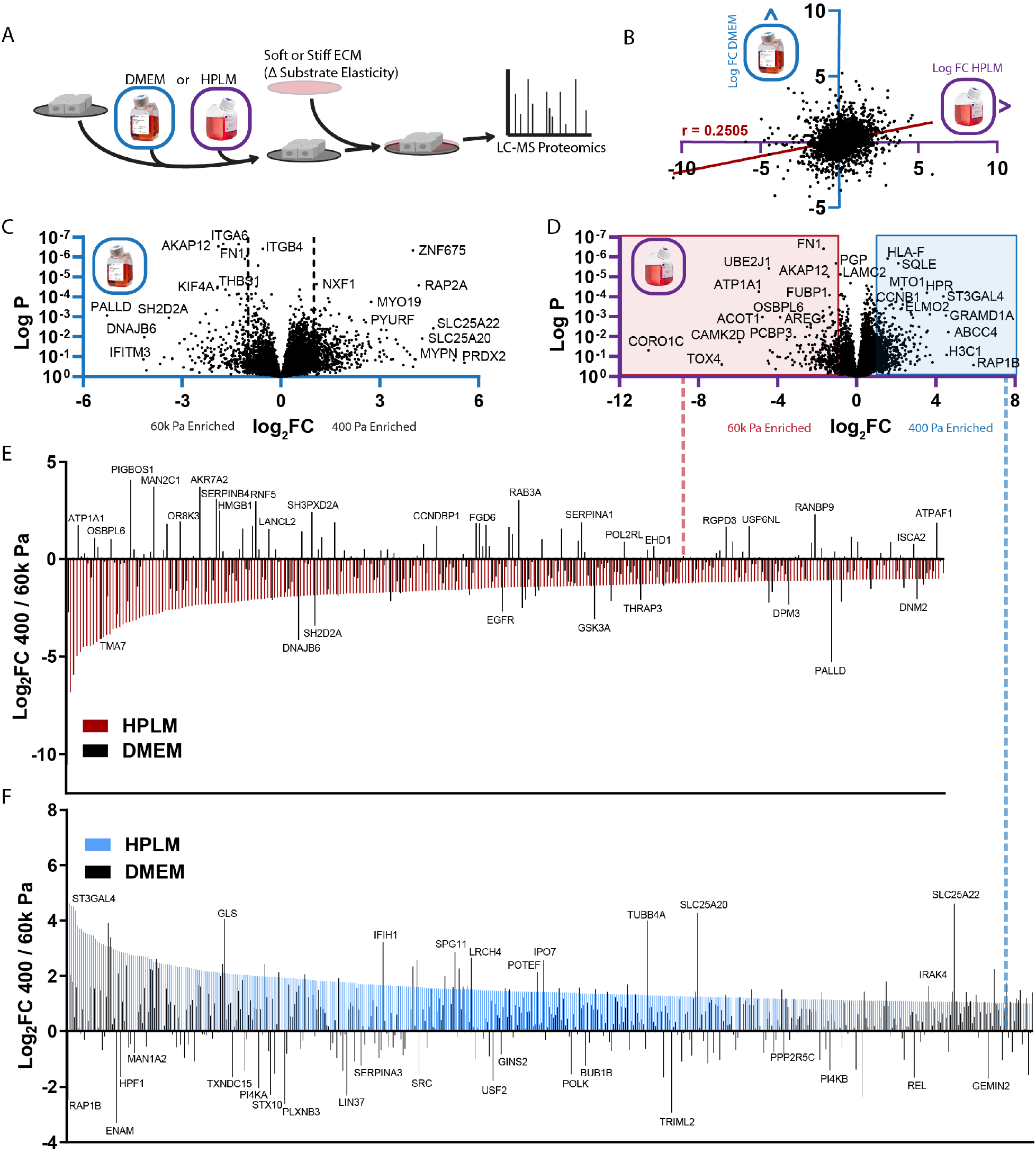
Culture medium dictates proteomic mechanoresponse A. Graphical representation of experimental design related to Figure 1B-F. (n = 3 biological replicates) B. Comparison of protein abundance between MCF10A cells cultured on 400 Pa (soft) vs 60k (stiff), either in HPLM (X-axis) or DMEM (Y-axis). Each point represents a single peptide’s fold change (400/60k) in a given medium, Pearson’s r-value = 0.2505, P value <0.0001 (two-tailed). C. Volcano plot depicting relative abundance of proteins in MCF10A cells cultured on 400 Pa vs 60k in DMEM (fold change, 400/60k). D. Volcano plot depicting relative abundance of proteins in MCF10A cells cultured on 400 Pa vs 60k in HPLM (fold change, 400/60k). E. Abundance rank order plot of proteins two fold enriched in MCF10A cells cultured on 60k Pa vs 400 Pa in HPLM (red, related to red box in D), with the same comparison in DMEM (black) superimposed (related to B). F. Abundance rank order plot of proteins two fold enriched in MCF10A cells cultured on 400 Pa 60k Pa in HPLM (blue, related to red box in D), with the same comparison in DMEM (black) superimposed (related to B).

To determine which cellular processes respond to mechanical cues, we performed Gene Ontology (GO) analysis on the differentially expressed proteins. The top GO category enriched in cells cultured in stiff HPLM microenvironments was “R-HSA-2262752: Cellular response to stress” which associates glucose metabolism, mitochondrial reactive oxygen species (ROS), DNA damage, and heat shock factor 1 (HSF1)-mediated adaptations to protein folding stress in multiple cellular compartments (Fig. S1G). Further analysis suggested that transcriptional mediators of mechanoresponses are Poly [ADP-ribose] polymerase 1 (PARP1), Krueppel-like factor 10 (KLF10), Hypoxia Inducible Factor 1 Subunit Alpha (HIF1A), Nuclear factor NF-kappa-B (RELA/NFKB1), and HSF1 (Fig. S1H) which regulates mitochondrial oxidation of glucose (*9*) and cytoskeletal integrity (*28*).

### ECM stiffness enhances metabolic support of glycosylation

Because concentrations of metabolic substrates influence its metabolic fate and drive cell-intrinsic mechanoresponses (*9*), we used mass spectrometry to measure metabolite levels of cells cultured in DMEM or HPLM with and without physiologically relevant hyperglycemia [15 mM] on a gradient of adhesive substrate stiffnesses (400, 6k, and 60k Pa) (Fig 2). As expected, principal component analysis revealed that medium composition (PC1) was the most deterministic factor of intracellular metabolite profiles (Fig. S2A), but we could also detect synergy between hyperglycemia and adhesion substrate stiffness (PC2). Examining medium-dependent mechanoresponse-induced metabolite changes, we noticed that stiff and hyperglycemic DMEM conditions biased cells to accumulate a number of nucleotides, which significantly altered the ATP:AMP ratio (Fig. S2B), whereas NADH levels increased with adhesive substrate stiffness in a medium-independent fashion (Fig. S2C). Stress-induced PARP1 activity (Fig. S1F) has been correlated with alterations in NAD(P)H photochemistry (*29*). Therefore, we used phasor-FLIM (fluorescence lifetime imaging microscopy) to monitor NAD(P)H dynamics. Surprisingly, we found that NAD(P)H photochemistry was only sensitive to adhesive substrate stiffness when cells were cultured in HPLM, but not DMEM (Fig. S2D-F).

**Figure 2:**
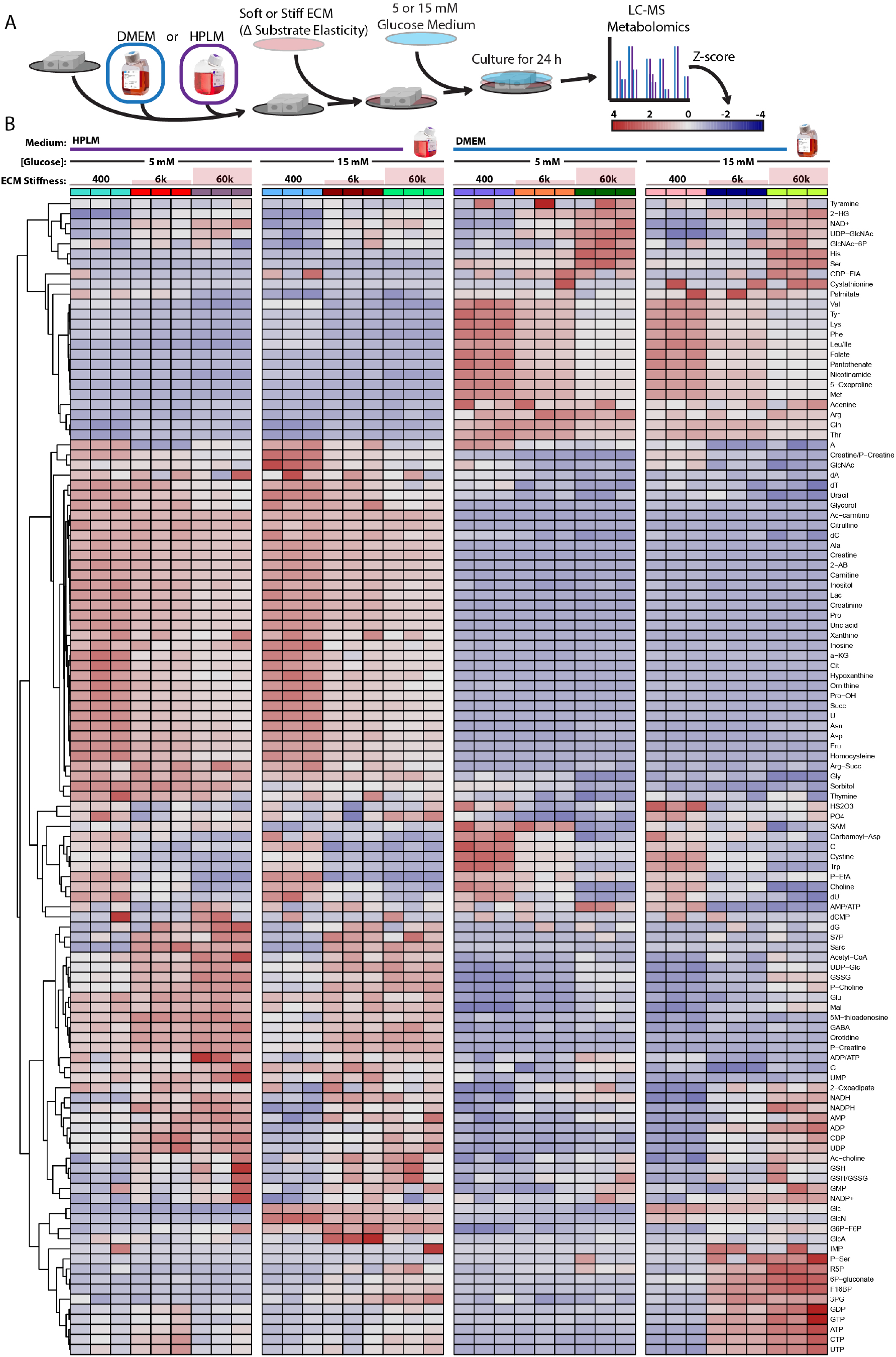
Culture medium dictates metabolomic mechanoresponse A. Graphical representation of experimental design related to Figure 2B. (n = 3 biological replicates) B. Heatmap depicting the relative abundance (z-score) of metabolites between cells cultured on 400 Pa, 6k Pa, or 60k Pa in HPLM or DMEM with 5 or 15 mM glucose. (n = 3 biological replicates, Z-score color scale depicted in A)

Examining the regulatory patterns of the composite metabolomic and proteomic effects of mechanoresponses in DMEM and HPLM (*30*), we found that NAD(P)H photochemistry and stress response signatures of cells cultured in stiff HPLM microenvironments were associated with increased glutathione oxidation, suggesting that stiff microenvironments elicit more oxidative stress and PARP1 activity (Fig. S1G-H and S3A-C). We also identified medium-independent mechanosensitive metabolic responses including the metabolism of arginine to ornithine (Fig. S3D) (*10*) and creatinine to phosphocreatine (Fig. S3E) (*5*). Interestingly, cells cultured in soft DMEM microenvironments may have increased synthesis of N-acetylglucosamine 6-phosphate (GlcNAc-6P) from endogenous N-acetylglucosamine (GlcNAc) because N-acetylglucosamine kinase (NAGK) is enriched (2.3 fold) (Fig. S3F-G). NAGK is a critical enzyme for the salvage/recycling of GlcNAc from lysosomal degradation of glycoconjugates (glycan conjugated proteins or lipids) when exogenous nutrients are limiting cellular objectives (*31*). NAGK levels were insensitive to adhesive substrate stiffness in HPLM, whereas hyperglycemia increased levels of GlcNAc-6P and abolished the mechanoregulation of NAGK suggesting that exogenous glucose could support GlcNAc demands via the hexosoamine biosynthetic pathway that uses glucose as a substrate to synthesize uridine diphosphate N-acetylglucosamine (UDP-GlcNAc).

UDP-GlcNAc and UDP-glucose (UDP-Glc) concentrations each increased with adhesion substrate stiffness in both media (Fig. 3A). While the relative concentration of UDP-GlcNAc did not differ between the two media, UDP-Glc levels were two-fold lower in DMEM. These UDP-charged sugars can be used to support the secretory pathway or glycocalyx synthesis, a carbohydrate-based pericellular matrix, by serving as a substrate for glycosylation in the endoplasmic reticulum and golgi apparatus (*32*). We directly measured isotopic glucose (^13^C) incorporation into UDP-GlcNAc and UDP-Glc in cells cultured on varied adhesion substrate stiffnesses in DMEM or HPLM. Fractional labeling patterns for UDP-Glc (greater) and UDP-GlcNAc (lower) dramatically differed for cells in DMEM versus HPLM (Fig. 3B). We reason that reduced incorporation of ^13^C into UDP-GlcNAc in DMEM-cultured cells could be explained by the fact that cells in these conditions favored NAGK-mediated salvage to generate UDP-GlcNAc, which was suggested by analysis of our multi-omics datasets. An explanation for why we detected a greater proportion of ^13^C-Glc without an increase in total UDP-Glc in cells cultured in DMEM could be that UDP-Glc was both synthesized and consumed at a greater rate.

**Figure 3:**
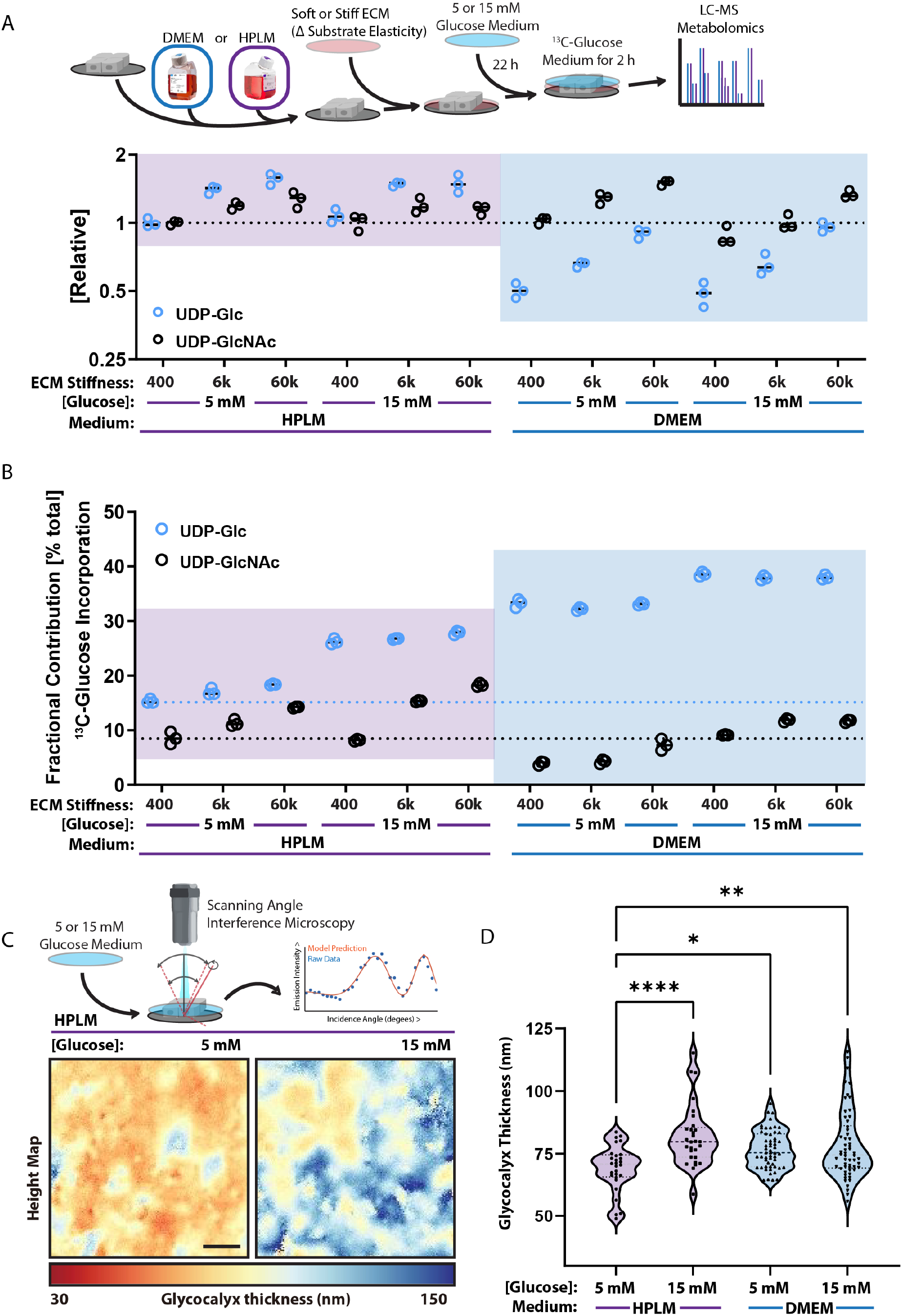
Microenvironmental properties dictate glycosylation and glycocalyx dynamics A. Graphical representation of experimental design related to Figure 3A-B and plot of relative abundance of UDP-Glc and UDP-GlcNAc between MCF10A cells cultured on 400 Pa, 6k Pa, or 60k Pa in HPLM or DMEM with 5 or 15 mM glucose. (n = 3 biological replicates) B. Plot depicting the percent of the total UDP-Glc and UDP-GlcNAc pool containing ^13^C derived from exogenous ^13^C_6_-glucose (fractional contribution) in cells cultured on 400 Pa, 6k Pa, or 60k Pa in HPLM or DMEM with 5 or 15 mM glucose. (n = 3 biological replicates) C. Representative scanning angle interference microscopy (SAIM) heightmaps of glycocalyx thickness of MCF10A cells in HPLM with or without 15 mM glucose. D. SAIM-based quantification of glycocalyx thickness of MCF10A cells in HPLM or DMEM with and without 15 mM glucose, results are the mean ± S.D. of at least 13 cells per condition (repeated 3 separate times with similar effects).

One expected biosynthetic output from UDP-Glc in epithelial and endothelial cells is the synthesis of the glycoconjugates used to form the glycocalyx (*14*, *33*, *34*), which has been linked to mechanosignaling and cancer progression in stiff and fibrotic tumors (*11*, *32*, *35*). To assay the glycocalyx abundance in these cells, we used scanning angle interference microscopy (SAIM), an axial localization microscopy technique with nanoscale precision (*36*, *37*). SAIM revealed that cells cultured in DMEM had a thicker glycocalyx than cells cultured in HPLM (Fig. 3C-D). Moreover, hyperglycemia was sufficient to increase the thickness of the glycocalyx in cells cultured in HPLM but not DMEM. Effects of hyperglycemia on the glycocalyx also corresponded to greater fractional labeling of UDP-Glc with ^13^C from glucose without an increase in the total abundance of UDP-Glc. These observations would fit a condition in which a greater proportion of newly synthesized UDP-Glc/GlcNAc was immediately incorporated into glycoconjugate synthesis and cell surface accumulation.

### Microenvironmental control of glycocalyx thickness, composition, and topology

To test if medium-dependent metabolic responses were associated with the formation of different glycoconjugates, we utilized lectin-based microarrays to survey the relative abundance of specific glycoconjugates formed (Fig. 4A) (*38*–*41*). Lectins are carbohydrate binding proteins that can detect specific glycan epitopes, distinguishing between epimeric and isomeric structures that have identical masses but different stereo and regiochemistry (e.g. Galβ1,3GalNAc and Galβ1,4GlcNAc) (*42*). Dual color lectin microarray analysis rapidly informs on combinatorial structures in the glycome that have biological significance in evolution/physiology. We found stark separation between the lectin-substrate signatures of cells cultured in DMEM and HPLM. Cells cultured in DMEM were enriched in glycoconjugates containing high mannose (GRFT, BanLec), oligo mannose (GNA), and terminal GalNAc (VVA, AIA). A potential signal for core fucose (PSA, LcH) was also observed, however these lectins have cross-specificity for high mannose. This indicates a less complex N-glycan set in DMEM cultured cells. In contrast, cells cultured in HPLM were enriched with glycoconjugates containing polyLacNAc (DSA, GSL-II), indicative of complex, branched N-glycans, and α2 Fucose (SNA-II, TJA-II), associated with Type 3/4 blood group H. Physiological hyperglycemia, which increased the thickness of the glycocalyx in HPLM but not DMEM (Fig. 3C-D), increased the abundance of α1,2 fucosylated epitopes in HPLM (PTA, Lewis Y) and DMEM (UEA) (Fig. 4B) and an increase in core 1/3 *O*-glycans, commonly found on mucins (AIA, MPA*). Glycan diversity and abundance can have significant impacts on cellular physiology (*33*), which makes the effects of culture medium a startling revelation.

**Figure 4:**
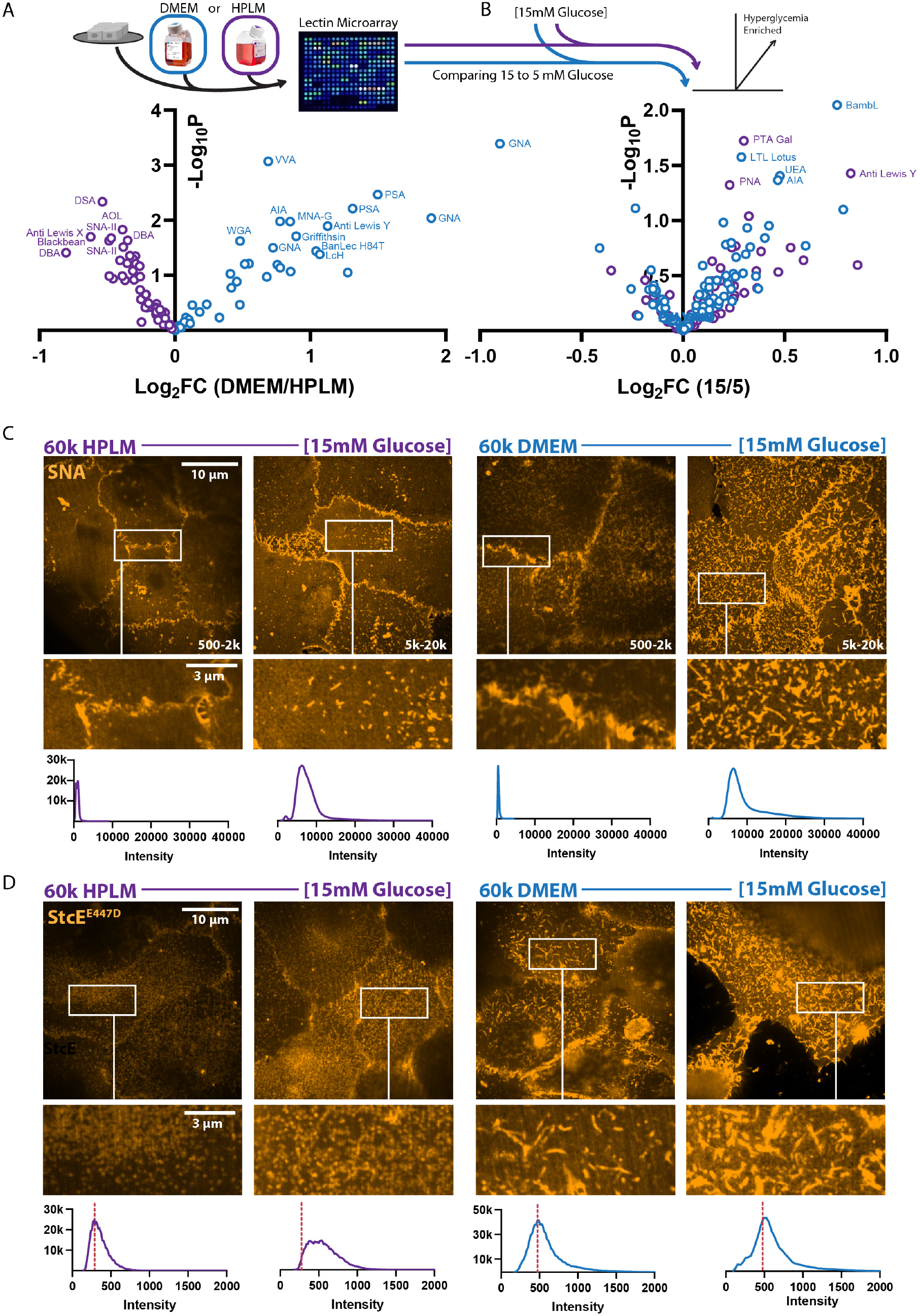
Microenvironmental properties dictate the abundance, topology, and composition of glycans that can comprise the cellular glycocalyx A. Volcano plot depicting relative abundance of glycans from MCF10A cells cultured in HPLM or DMEM, detected via lectin microarray. (n = 4 biological replicates) B. Volcano plot depicting relative abundance of glycans from MCF10A cells cultured in HPLM with 5 or 15 mM glucose, detected via lectin microarray. (n = 4 biological replicates) C. Representative super resolution via optical reassignment (SoRa) confocal microscopy of SNA staining of unpermeabilized MCF10A cells cultured on 60k Pa in HPLM or DMEM with or without 15 mM glucose. Associated fluorescent intensity histogram and intensity range of each image indicated in figure. D. Representative SoRa confocal microscopy of StcE^E447D^ staining of unpermeabilized MCF10A cells cultured on 60k Pa in HPLM or DMEM with or without 15 mM glucose. Associated fluorescent intensity histogram shown in figure.

While lectin microarrays provide a wealth of biochemical information, they do not clarify the sub-cellular localization of glycoconjugates (i.e., specific organelles or the cell surface). Because we wanted to understand glycocalyx dynamics, we used lectin staining of unpermeabilized cells with super resolution confocal microscopy to assess the abundance and topology of cell surface glycoconjugates. To this end, we used *Sambucus Nigra* lectin (SNA), which is specific to a α2,6-sialic acids found on complex *N*-glycans, for our initial analysis. We observed a ∼ 10-fold increase in fluorescence intensity in response to hyperglycemia with a punctate distribution across the cell surface and enrichment at cell-cell contact sites in HPLM (Fig. 4C). We observed a similar fluorescence intensity increase for hyperglycemia in DMEM cultured cells, but topologically the SNA staining showed ridge-like projections with irregular but interconnected vertices (Fig. 4C), resembling a Turing pattern.

Ridge-like glycocalyx topologies and projections may reflect a higher density of heavily glycosylated high molecular weight serine/threonine-linked glycoconjugates (e.g., mucins) (*43*). To determine if these cells contained greater concentrations or heterogeneous distributions of mucin-domain containing proteins on the cell surface we used a bacterial O-glycoprotease (StcE) with an enzyme-inactivating point mutation (StcE^E447D^) that retains affinity for mucin-domains of glycoproteins, which are heavily *O*-glycosylated (*44*, *45*). StcE^E447D^ staining of cells cultured in HPLM revealed a regular distribution of puncta across the cell surface, which increased in intensity in response to hyperglycemia (Fig. 4D). StcE^E447D^ staining of cells cultured in DMEM showed the same ridge-like projections with irregular but interconnected vertices we had observed with SNA staining, but hyperglycemia did not change the staining intensity on the cell surface. This is consistent with the increase in *O*-glycans observed by lectin microarray in HPLM, but not DMEM with hyperglycemia.

To measure the abundance of putative protein carriers of the SNA and StcE^E447D^ detected glycosylation we used mass spectrometry-based proteomics of cells cultured in soft and stiff microenvironments with and without hyperglycemia in DMEM or HPLM-based media (Fig. S4A). We found that the biochemical and mechanical properties of the microenvironment license differential responses to hyperglycemia. Paradoxically, these results revealed that hyperglycemia reduced the abundance of the immunomodulatory glycoprotein CD58 in cells cultured in stiff HPLM microenvironments (*46*–*48*). We observed a similar effect in DMEM microenvironments for CD46, another immunomodulatory glycoprotein. This may be because we designed our experiments to measure the abundance of glycoproteins without accounting for the fact that many of these glycoconjugates are also secreted and would not be quantified in the membrane associated proteome of in vitro cultures if they were secreted at a greater rate (*49*). Secretory programming would logically provide a means to remodel or construct the cellular glycocalyx, since newly synthesized glycoconjugates use secretory programs to traffic to the plasma membrane.

When we examined the ontology of the proteins that showed increased expression in response to hyperglycemia in soft or stiff microenvironments (Fig. S4B-C), we were confronted with the fact that the top category in soft microenvironments was indeed vesicular-mediated transport, which includes many secreted or secretion-facilitating proteins. StcE^E447D^ staining showed that cells cultured in soft HPLM microenvironments were coated in a regular pattern of circular/spherical structures, whereas cells in soft DMEM microenvironments displayed ridge-like projections with irregular but interconnected vertices (Fig. S4D). This topological distribution of StcE^E447D^ staining of cells cultured in soft HPLM environments with hyperglycemia led us to suspect that, as the proteomics suggested, hyperglycemia promotes the secretion of vesicles decorated with CD58 or other glycoproteins.

### HSF1 mediates hyperglycemia induced glycocalyx construction

Since hyperglycemia was sufficient to increase the glycocalyx thickness of cells cultured in HPLM, we decided to test if we could identify positive regulators of glycocalyx construction by examining which proteins become more abundant in response to hyperglycemia. Promotor analysis of the hyperglycemia-enriched proteins revealed a number of putative transcriptional mediators, such as PPARGC1A, CREB, YY1, HSF1/2, ARNT2, USF and an unassigned motif (ACTAYRNNNCCCR) were correlated with increased glycocalyx abundance (Fig. 5A). When we separated these putative mediators by their effects on cells cultured in soft or stiff microenvironments, only proteins with the unassigned motif (ACTAYRNNNCCCR) were more abundant because of hyperglycemia (*50*). However, the ACTAYRNNNCCR-associated proteins induced by hyperglycemia were completely inconsistent and unrelated between soft (Glycerolipid Biosynthesis) and stiff microenvironments (Cellular Response to DNA Damage Stimulus and Signaling by Rho GTPases). We noticed that hyperglycemia caused cells in soft microenvironments to enrich for many of the same proteins predicted to be downstream of PPARGC1A, HSF1, and HSF2, whereas cells in stiff microenvironments enriched for a more broad “HSF” assignment. HSF1 is a transcription factor associated with canonical (heat shock) and non-canonical responses (*51*) that modulate glucose metabolism to support stress remediation programs (*9*, *52*–*54*). Because of these associations, we assayed isotopic glucose (^13^C) incorporation into UDP-GlcNAc and UDP-Glc in cells cultured in HPLM with or without an HSF1 inhibitor (KRIBB11) and found that HSF1 inhibition significantly reduced the fractional contribution of ^13^C-glucose derived carbons into UDP-Glc and UDP-GlcNAc (Fig. 5B).

**Figure 5:**
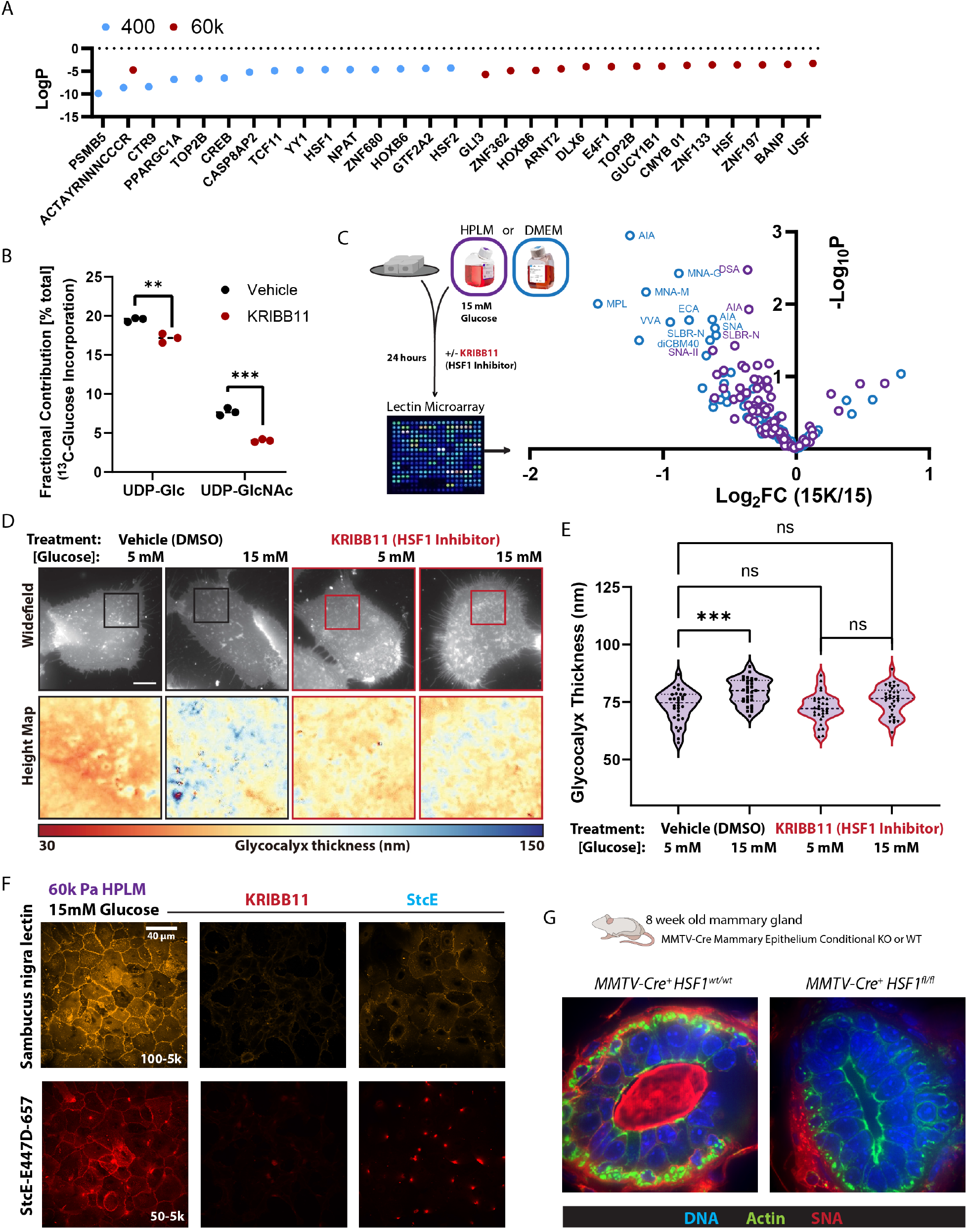
HSF1 mediates hyperglycemia induced glycocalyx construction A. Transcriptional Regulatory Relationships Unraveled by Sentence-based Text mining (TRRUST) based detection of transcription factors associated with the proteins significantly enriched in response to 15 mM glucose in HPLM on 400 Pa or 60k Pa. (n = 3 biological replicates LC-MS based proteomics) B. Fractional contribution of ^13^C_6_-glucose to UDP-Glc and UDP-GlcNAc in cells cultured in HPLM +/- KRIBB11 [2 µM]. (n = 3 biological replicates) C. Volcano plot depicting relative abundance of glycans from MCF10A cells cultured in HPLM with 15 mM glucose +/- KRIBB11 [2 µM], detected via lectin microarray. (n = 4 biological replicates) D. Representative SAIM-based heightmaps of glycocalyx thickness of MCF10A cells in HPLM with 15 mM glucose +/- KRIBB11 [2 µM]. E. SAIM-based quantification of glycocalyx thickness of MCF10A cells in HPLM with 15 mM glucose +/- KRIBB11 [2 µM], results are the mean ± S.D. of at least 13 cells per condition (repeated 3 separate times with similar effects). F. Representative confocal microscopy of SNA and StcE^E447D^ staining of unpermeabilized MCF10A cells cultured in HPLM with 15 mM glucose +/- KRIBB11 [2 µM] or StcE [2 µM]. G. Representative confocal microscopy of SNA (red), phalloidin (green), and DAPI (blue) staining of 8 week old murine mammary gland acini, from MMTV-Cre^+^ mice and *MMTV-Cre^+^Hsf1^fl/fl^* mice. (n = at least 5 acini were examined from 3 WT and 5 KO mice, all KO acini had no detectable SNA).

To determine if the reduced metabolism of exogenous glucose into UDP-Glc and UDP-GlcNAc caused by HSF1 inhibition was sufficient to suppress hyperglycemia induced glycocalyx construction, we assayed cells cultured in hyperglycemia with and without HSF1 inhibition with a lectin microarray. HSF1 inhibition prevented hyperglycemia from increasing the abundance of the [glucose]-responsive glycans (Fig. 5C) and reduced the abundance of sialoglycans of cells in normal [glucose] concentrations (Fig. S5A). Using SAIM, we were able to see that HSF1 inhibition was sufficient to block the thickening of the glycocalyx that occurred in response to hyperglycemia (Fig. 5D-E). HSF1 inhibition did not reduce the thickness of the glycocalyx below the height of cells cultured in HPLM, which could indicate that the HSF1 program is suited to regulate and route the metabolism of glucose beyond a baseline concentration threshold (i.e., which could explain why we detected an HSF1 driven response to hyperglycemia). This is consistent with the fact that HSF1 regulates the expression of many heat shock proteins (HSPs) that were also discovered because they became more abundant in response to glucose, and thus an alias for many HSPs are also Glucose Regulated Proteins (GRP) (*55*).

Staining of mammary epithelial cells with SNA and StcE^E447D^ revealed that HSF1 inhibition reduced the abundance of cell surface sialic acid and mucin-domain containing glycoconjugates more than treatment with a high concentration [2 µM] of the bacterial *O*-glycoprotease, StcE (*56*), which cleaves proteins conjugated with mucin-type *O*-glycosylation (Fig. 5F and S5B-C). To verify the relationship between HSF1 and sialic acid abundance *in vivo*, we generated a mouse model in which HSF1 was conditionally deleted in mammary epithelial cells (*Hsf1* floxed alleles (*57*) crossed with the MMTV-Cre, line D). In *MMTV-Hsf1KO* mammary tissues, we found no SNA staining of the lumens or cell-cell junctions of the mammary epithelial acini, consistent with the loss of this epitope observed by lectin microarray. In contrast, in the surrounding mammary adipose tissue SNA staining was unaffected (Fig. 5G and S5D). Overall, these results connect physiologically relevant hyperglycemia to HSF1 regulated glycocalyx construction on the surface of mammary epithelial cells *in vitro* and *in vivo*.

### Glycocalyx composition and topology affect Natural Killer cell immunogenicity

We next sought to test how microenvironment-disposed mechanoresponses affect the composition and topology of the cellular glycocalyx since critical enzymes (e.g., ST3GAL4) and metabolite fluxes (e.g., UDP-GlcNAc) were differentially abundant as a result of adhesion substrate stiffness. Using lectin microarrays, we found that α-2,3 -sialic acid on Type II LacNAc (Neu5Acα1-2Galβ1-4GlcNAc, detected by SLBR-N) was significantly decreased in response to stiffness (Fig. 6A) (*58*). N-acetylneuraminic acid (Neu5Ac) is the predominate sialic acid species observed in human cells. This data correlates well with the loss of ST3Gal4 observed in the proteomics. The ST3Gal4 enzyme transfers Neu5Ac onto the 3-position of galactose of Type II LacNAc motifs to create the α-2,3 sialylated epitope, which influences outcomes of immunogenic receptor signaling (*18*, *59*). De novo biosynthesis of Neu5Ac occurs via UDP-GlcNAc 2-epimerase/ManNAc kinase (GNE/MNK), N-acetylneuraminate synthase (NANS), and Neu5Ac9-P-phosphatase (NANP) mediated metabolism of UDP-GlcNAc/ManNAc to ManNAc-6-phosphate to Neu5Ac-9-phosphate to Neu5Ac. For Neu5Ac to be linked to a glycan, it must be charged with cytidine monophosphate (CMP) in the nucleus by the CMP sialic acid synthase (CMAS). The activated sugar, CMP-Neu5Ac, exits the nucleus and is transported into the golgi apparatus for incorporation into glycoconjugates via a number of sialyltransferases (e.g., ST3Gal4) (*62*–*64*).

**Figure 6:**
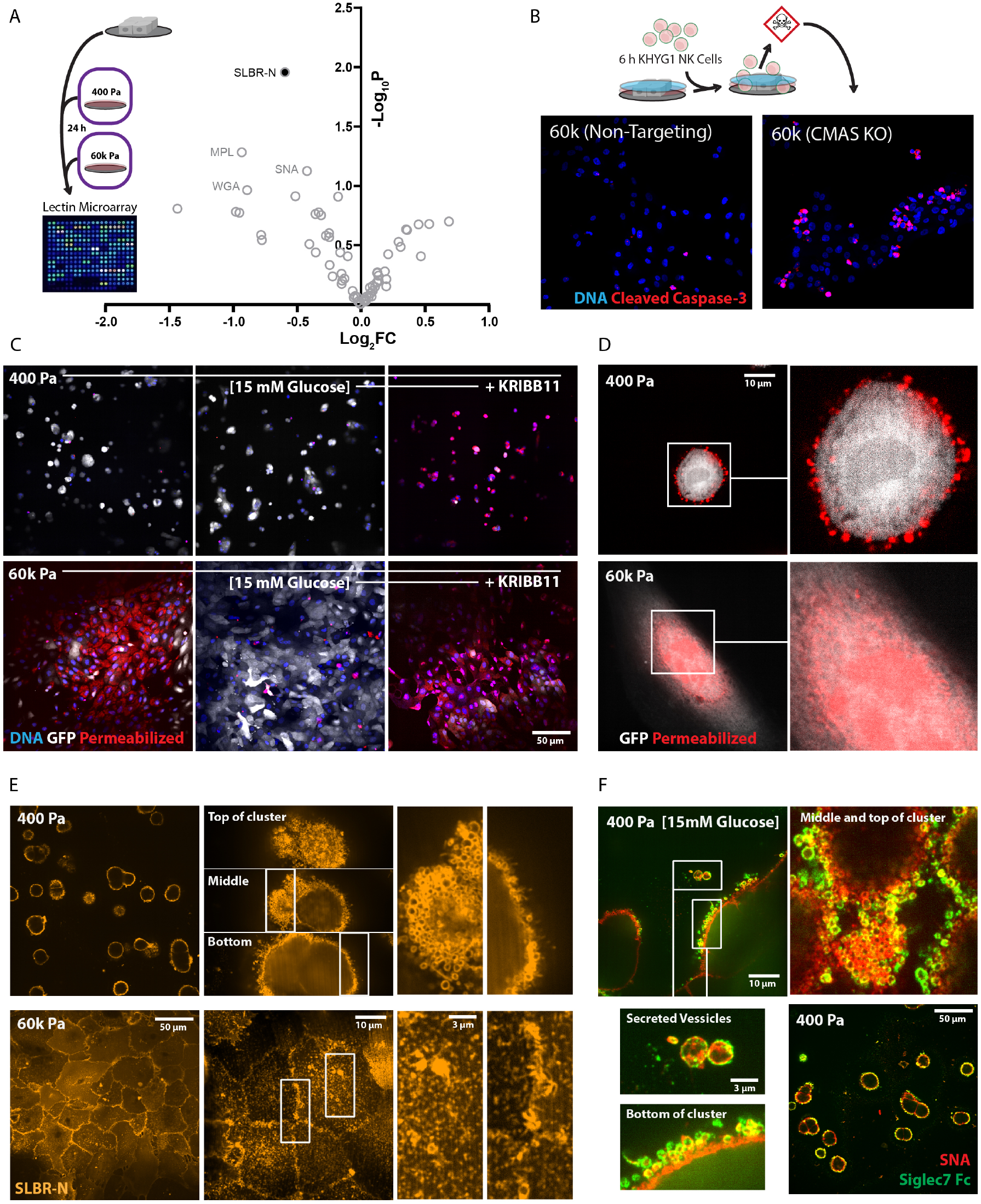
The physical properties of the microenvironment affect Natural Killer cell immunogenicity via glycocalyx composition and topology A. Volcano plot depicting relative abundance of glycans from MCF10A cells cultured in HPLM on 400 Pa or 60k Pa, detected via lectin microarray. (n = 4 biological replicates) B. Representative confocal microscopy of cleaved caspase 3 (red) and DAPI (blue) staining of non-targeting WT controls and CMAS KO MCF10A cells after 6 h of co-culture with equal numbers of KHYG1 NK cells in HPLM. (repeated twice with similar results) C. Representative confocal microscopy of propidium iodide homodimer (red) and DAPI (blue) staining of cytosolic-GFP (white) expressing MCF10A cells on 400 Pa or 60k Pa after 24 h of co-culture with equal numbers of KHYG1 NK cells in 5 mM glucose HPLM. Treatments indicated (+/-15 mM glucose or 15 mM glucose + KRIBB11 [2 µM]) occurred for the 24 h prior to the addition of equal number of KHYG1 cells to the MCF10A cultures in HPLM with 5 mM glucose and no KRIBB11. (repeated three times with similar results) D. Representative SoRa confocal microscopy of propidium iodide homodimer (red) staining of a cytosolic-GFP (white) expressing MCF10A cell on 400 Pa or 60k Pa after 24 h of co-culture with equal numbers of KHYG1 NK cells in HPLM. E. Representative confocal (left panel) and SoRa confocal microscopy (right panel and enhancements) of SLBR-N staining of MCF10A cells on 400 Pa or 60k Pa in HPLM. F. Representative confocal (left panel) and SoRa confocal microscopy (right panel and enhancements) of SLBR-N staining of MCF10A cells on 400 Pa or 60k Pa in HPLM. G. Representative confocal (bottom right) and SoRa confocal microscopy (top and expansions) of recombinant Siglec7 Fc (green) and SNA staining of MCF10A cells on 400 Pa or 60k Pa in HPLM.

To test if the sialoglycans detected by SNA and SLBR-N in cells cultured in stiff HPLM microenvironments influence immunogenicity of the glycocalyx, we engineered a CMAS knockout human mammary epithelial cell line (CMAS KO). CMAS deletion did not affect the total cell surface mucin content (i.e., no difference in StcEE^447D^ staining), but did increase truncated mucin-type O-glycans measured by *Bacteroides thetaiotaomicron* derived metalloproteinase (BT4244) with an inactivating point mutation (BT4244^E575A^) staining (*44*). BT4244^E575A^ preferentially binds truncated core 1 O-glycans (e.g., T (Gal-GalNAc) and Tn (GalNAc) antigens) that lack a terminal sialic acid, the predicted outcome of a CMAS deletion (Fig. S6A). Sialoglycan interactions with receptors like Siglec7, which binds α-2,3-sialosides, affect Natural Killer (NK) cell activation and killing-efficiency (*19*), so we used a human NK cell line (KHYG1 (*65*)) to test if CMAS KO cells were more sensitive to NK cell mediated killing. As expected, in stiff HPLM microenvironments global ablation of sialoglycoconjugates by CMAS KO caused an increase of the number cells stained positive for cleaved caspase 3 (apoptotic) after 6 hours of co-culture with a matched concentration of KHYG1 cells (Fig. 6B). These data confirm that cells with greater cell surface sialyation are more resistant to NK cell mediated killing (*37*, *66*).

Because we found that hyperglycemia increased the amount of cell surface sialyation, and that HSF1 inhibition was sufficient to prevent this, we assayed NK cell-mediated killing of mammary epithelial cells previously cultured in hyperglycemia with and without KRIBB11 on soft or stiff microenvironments (Fig. 6C). As predicted, cells cultured in stiff HPLM microenvironments were less sensitive to NK cell-mediated killing if they were exposed to hyperglycemia prior to the 24-hour NK cell challenge. HSF1 inhibition was sufficient to abolish the NK cell resistance conferred by hyperglycemia in stiff microenvironments. Surprisingly, we found that hyperglycemia had little effect on the viability of cells cultured in soft HPLM microenvironments, not because hyperglycemia failed to increase NK cell resistance, but because the cells in soft microenvironments were innately resistant to NK cell-mediated killing, regardless of the glucose concentration. However, HSF-1 was sufficient to overcome this resistance, in line with its impact on the glycome.

NK cells kill target cells by first permeabilizing target cells with perforin, so that they are able to deliver granzymes, which activate apoptotic pathways in the target cell once they reach the cytosol (*65*). The fact that epithelial cells cultured in stiff microenvironments were more susceptible to NK cell-mediated apoptosis could be due to the stress responses (*66*) activated by cells in those microenvironments (Fig. S3D). However, when we examined what NK cells had permeabilized, we found that cells cultured in soft microenvironments had been permeabilized by the NK cells, just not their cytosol (Fig. 6D). In assessing what was permeabilized, we found that the plasma membrane of these cells contained puncta that were permeabilized, but these puncta were distinct from the cytosolic-GFP expressed in the cells used for these experiments. In contrast, cells cultured on stiff microenvironments were permeabilized throughout their cytosol and nucleus.

It was surprising to observe the permeabilized plasma membrane puncta in cells cultured in soft HPLM microenvironments that survived the NK cell challenge, but we had also observed with SNA and StcE^E447D^ staining that cells in soft microenvironments appeared to be coated in a regular pattern of circular/spherical bleb- or vesicle-like structures (Fig. S4D). We then checked if SLBR-N staining revealed similar topologies (Fig. 6E). We observed that the extremities of cells in soft microenvironments were wrapped in sialic acid labeled bleb- or vesicle-like structures, analogous to a glycan bubble-wrap we refer to as a sialic acid bleb-sicle surface (SABS). In stark contrast, cells cultured in stiff microenvironments accumulated sialic acid in-between cell-cell junctions and displayed a regular pattern of SLBR-N stained puncta across the cell surface. We verified that these structures were also detected with both SNA and the Siglec7 Fc (Fig S6B), and found that these vesicles were secreted from the cell surface in response to hyperglycemia (Fig. 6F), which was predicted by our proteomics (Fig. S4C).

We hypothesized that sialic acid may be required for the formation of vesicular structures detectable with SNA and SLBR-N that formed in soft HPLM microenvironments. Testing this dependency presented a challenge, since the ability to visualize these structures was dependent on the glycan modification we sought to remove. However, wheat germ agglutinin (WGA) is a lectin that detects both GlcNAc and Neu5Ac terminated glycans. Using WGA and SNA, we were able to detect co- localized fluorescent signal of SABS (Fig S6C) in control cells. However, the SABS topology was absent from CMAS KO cells cultured in soft microenvironments indicated that sialic acid is a necessary component of these structures. Because these structures were mechanosensitive, we wanted to determine the adhesive substrate stiffness at which they would disappear. Using a range of stiffnesses that span age-associated changes of the elastic modulus (Fig. S6D-E) of the murine mammary gland, we found that above 1k Pa, the vesicular structures disappeared in favor of the puncta we observed in the microenvironments that recapitulated tumor elasticities.

ECM stiffness engenders cell spreading via cytoskeletal rearrangements supported by cell-ECM adhesions. To test if cytoskeletal dynamics affect the maintenance of SABS, we utilized an inducible Rho-associated protein kinase (ROCK) model (ROCK::ER) (*67*) that promotes cytoskeletal remodeling and spreading. ROCK::ER activation was sufficient to dissipate the bleb-sicle phenotype in favor of spreading of single cells and clusters (Fig. S7A). ROCK::ER activation also caused the accumulation of sialic acid within cell-cell junctions (Fig. S7A). Taken together, increased cell spreading in soft microenvironments is sufficient to recapitulate the cell surface sialic acid topology and distribution of cells cultured in stiff microenvironments. Conversely, human Embryonic Stem Cells (hESCs) have enhanced viability when treated with the ROCK inhibitor (Y27632) (*68*) and ROCK inhibition was sufficient to generate the SABS phenotype on hESC (Fig. S7B).

As spreading cells adjust their geometries and generate contractile forces via cell-ECM adhesions, intracellular water effluxes into the microenvironment through the semi-permeable plasma membrane (*69*, *70*). Similarly, hypertonic crystalloid or colloid solutions reduce intracellular water concentrations, and DMEM has a higher osmolality (320-355 mM/kg) than human plasma or extracellular fluid (275-299 mM/kg). The discrepancy between DMEM and plasma osmolality is largely due to differences in inorganic salt concentrations. To test if the salt composition of DMEM was sufficient to prevent SABS formation in soft microenvironments, we created a derivative of HPLM containing DMEM-defined inorganic salts (HPLM^DMEM^ ^Salt^), which had a similar osmolality to DMEM (Fig. 7A). MCF10A cells cultured in HPLM^DMEM^ ^Salt^ did not display SABS (Fig. 7B) despite similar levels of WGA surface staining (Fig. S7C). To verify that osmotic pressure was sufficient to dissipate the blebsicle surface phenotype, we treated cells with polyethylene glycol (PEG)-supplemented HPLM. PEG-mediated osmotic pressure increased glycocalyx thickness (Fig. S7D) and dissipated the SABS phenotype (Fig. 7C).

**Figure 7:**
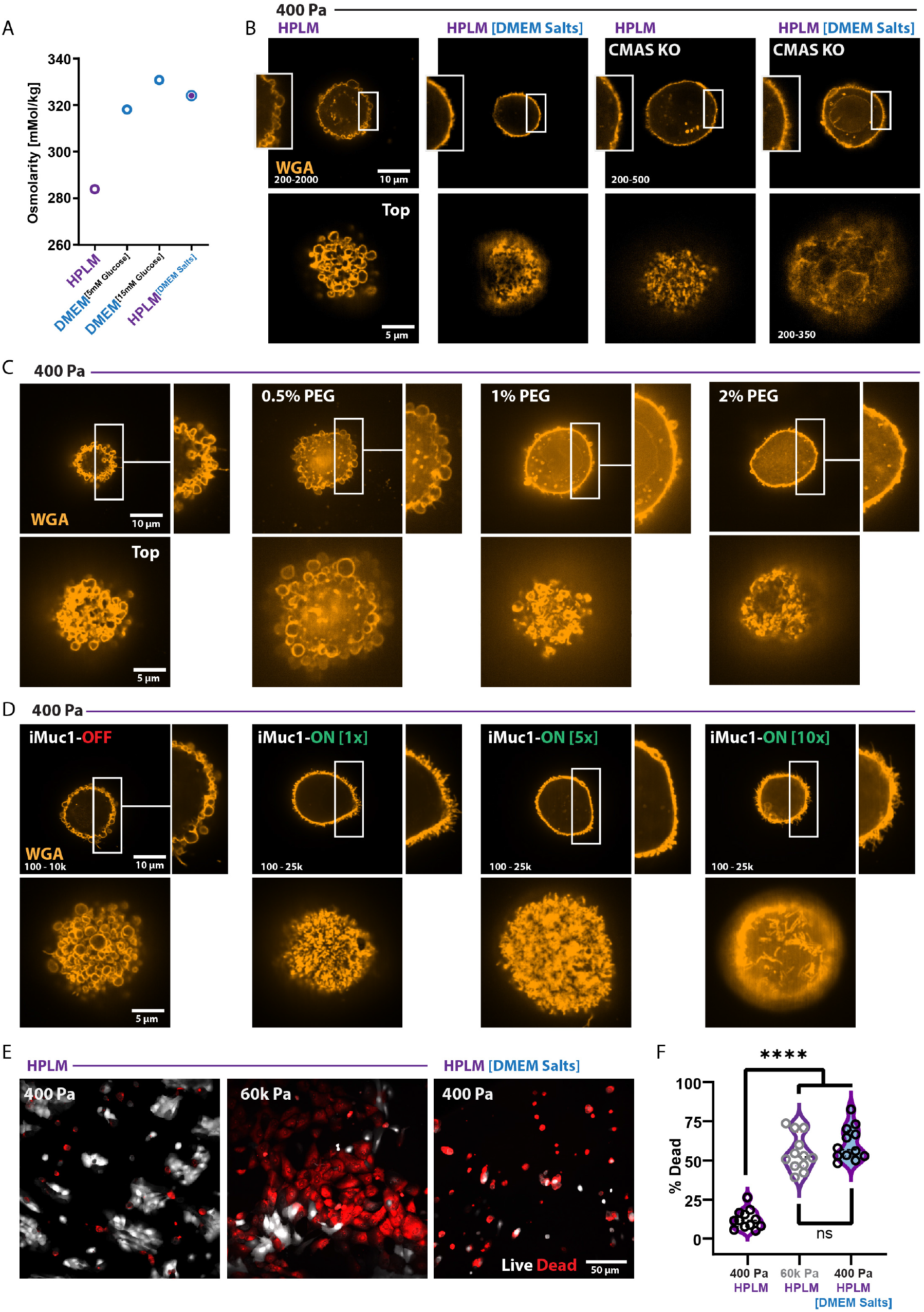
Microenvironmental regulation of glycocalyx topology is related to osmotic adjustments and dictates Natural Killer cell immunogenicity A. Osmolality of additive-free culture media. B. Representative SoRa confocal microscopy of WGA staining of unpermeabilized non-targeting WT controls and CMAS KO MCF10A cells on 400 Pa in HPLM or HPLM^DMEM Salt^. C. Representative SoRa confocal microscopy of WGA staining of unpermeabilized MCF10A cells on 400 Pa in HPLM with increasing concentrations of PEG-400 (v/v %) for 24 h. D. Representative SoRa confocal microscopy of WGA staining of unpermeabilized iMuc1-MCF10A cells on 400 Pa in HPLM (1x = 1 ng/mL, 5x = 5 ng/mL, 10x = 10 ng/mL Dox). E. Representative confocal microscopy of propidium iodide homodimer (red) and calcien-AM (white) staining of MCF10A cells on 400 Pa or 60k Pa after 24 h of co-culture with equal numbers of KHYG1 NK cells in HPLM or HPLM^DMEM^ ^Salt^. (repeated three times with similar results) F. Quantitation of effects shown in E from 10 field views randomly selected across 2 biological replicate experiments.

Because PEG-mediated osmotic pressure increased glycocalyx thickness, we sought to determine if increased glycocalyx thickness was sufficient to dissipate the SABS phenotype. To test the relationship between glycocalyx thickness and bleb formation we utilized inducible expression of Mucin 1 (iMUC1), a “bulky” cell surface glycoprotein that strongly influences net glycocalyx thickness (*37*). As predicted, increased glycoclayx thickness dissipated the SABS phenotype (Fig. 7D). Conversely, reducing glycocalyx thickness with mucinase treatment (StcE) was sufficient to generate small regions of SABS on every cell in stiff hyperglycemic microenvironments (Fig. S7E). Without sialic acid (CMAS KO), PEG-mediated osmotic pressure did not increase glycocalyx thickness because the cell membranes ruptured and all of the CMAS KO cells died (Fig. S7F-G). Overall, the data indicate that ECM stiffness and culture medium influence the topology of sialic acid glycoconjugates via interrelated changes in glycocalyx thickness and cytoplasmic pressure.

Now that we understood that osmotic pressure caused by the properties of the microenvironment was sufficient to dissipate the SABS phenotype, we decided to revisit the relationship between cell surface sialic acid topology and sensitivity to NK cell-mediated killing. MCF10A cells cultured in soft microenvironments displayed the SABS phenotype and were more resistant to NK cell-mediated death than the same cells cultured in stiff microenvironments, which did not display the SABS phenotype (Fig. 6C). MCF10A cells cultured in soft HPLM^DMEM^ ^Salt^ microenvironments did not display the SABS phenotype despite having similar spread area and sialic acid surface content. Co-culture of NK and MCF10A cells in soft and stiff microenvironments with HPLM or HPLM^DMEM^ ^Salt^ showed significant differences in MCF10A cell survival, whereby cells without the SABS phenotype were ∼4 fold more likely to be killed by NK cells (Fig. 7E-F). Together, the data indicate that physical and biochemical properties of the microenvironment play a crucial and intertwined role in regulating glycocalyx topology, which is a previously unknown modulator of immune surveillance.

## Discussion

In this work, we have shown that the interplay of microenvironmental factors can dictate metabolic phenotypes, proteome dynamics, and glycocalyx construction of mammary epithelial cells largely through HSF1-mediated programming. We found that the physical properties of the microenvironment influence immune surveillance via compositional and topological dynamics of the sialic acid-containing glycocalyx. Contrary to past assumptions of the inertness of culture medium and adhesion substrate elasticity, we demonstrate that the biochemical and physical properties of the cellular microenvironment can obfuscate the immunogenicity of the glycome (i.e., composition and structure), markedly alter cellular composition (e.g., proteome), and tune metabolic programming (e.g., metabolomics and fate mapping). Therefore, translation-focused pre-clinical studies, including efforts to augment anti-tumor immunity or drug cancer cell autonomous programming, should be mindful of the cellular context for the models used because cellular physiology and sensitivities are disposed through a dynamic interplay with the microenvironment (*71*, *72*).

Mechanoresponses induced by the stiff and fibrotic mammary tumor microenvironment are associated with tumor progression, metastasis, and HSF1-mediated metabolic adaptions. HSF1 is a crucial regulator of metastatic progression and here we show that HSF1 regulates the abundance, composition, and immunogenicity of the cell surface glycome. Our findings indicate that synergy between ECM stiffness and hyperglycemia may engender metastasis via metabolic programming necessary to construct an immune suppressive glycocalyx. Given our findings, we predict that the glycocalyx may confer immune evasive characteristics to tumor cells in primary and secondary sites through different mechanisms. In the stiff primary tumor, glycocalyx bulk and sialic acid abundance may shield tumor cells from immune surveillance. In soft secondary tumor sites (e.g., lung and brain), the compositional (e.g., α2,3 sialylation) and structural dynamics of the glycocalyx (bleb-like vesicles) may preclude immune cell-mediated killing by excluding apoptosis-inducing factors from the cytoplasm (*73*).

## Acknowledgments

We thank the following people: Chenkai Dai (NIH) for providing us with the *Hsf1^flox^* mouse model; James Koh (UCSF) for facilitating osmolality measurements; Torsten Wittmann (UCSF) and Kyle Marchuk (UCSF) for acquiring and exquisite maintenance of the SoRa microscope; Mallar Bhattacharya (UCSF) for constructive review of the manuscript and discussions about the broader context of these findings; UCSF Laboratory Animal Resource Center (LARC) staff for care and housing of our animals; Casey Beppler (UCSF) for assistance with CAR-T experiments which were informative but not included in this manuscript; and Elizabeth Willey (UCB) for assistance with an epic tissue harvest.

## Author Contributions (Contributor Roles Taxonomy, CRediT)

Conceptualization: K.M.T. Methodology: K.M.T, G.A.T., M.J.P., L.K.M., D.L.S., & J.tH.-S. Software: J.A.B., S.P., & A.E.Y.T.L. Investigation: K.M.T., S.P., G.A.T., A.L.R., F.P., A.E.Y.T.L, N.J.T., N.M.E.A., & J.tH.-S. Data Curation: K.M.T., A.L.R., L.K.M., & A.E.Y.T.L. Resources: J.R.C., E.L.P., D.J.S., R.W.S., J.R.C., K.T., C.R.B., & N.M.E.A. Formal Analysis: K.M.T., S.P., A.L.R., F.P., A.E.Y.T.L, N.J.T., & J.tH.-S. Funding acquisition: V.M.W., M.J.P., K.M.T., D.L.S., L.K.M., A.D., C.R.B., Project administration: K.M.T. Supervision: K.M.T., V.M.W., M.J.P., J.R.C., D.L.S., & L.K.M. Visualization: K.M.T. Writing – original draft: K.M.T. Writing – review & editing: K.M.T, J.R.C., S.P., G.A.T. E.L.P., N.M.R., D.L.S., A.L.R, D.J.S., F.P., A.E.Y.T.L,, N.M.T., L.K.M., & V.M.M.

## Data and materials availability

All data are available in the main text or the supplementary materials and multiomics datasets will be available on the interwebz with a GUI explorer upon publication.

## Materials and Methods

### Human Cell Culture

MCF10A (ATCC) were originally passaged in [5 mM] glucose DMEM:F12 (1:1 mixture of F:12 [10mM] glucose and [0 mM] glucose DMEM) (Gibco, 11765054 and 11966025) supplemented with 5% Horse Serum (HS) (Gibco, 16050-122), 20 ng/mL epidermal growth factor (Peprotech), 10 µg/mL insulin (Sigma), 0.5 µg/mL hydrocortisone (Sigma), 100 ng/mL cholera toxin (Sigma, C8052-2MG), and 1x penicillin/streptomycin (Gibco). For experiments comparing HPLM to DMEM, a biological replicate cryopreserved stock of MCF10A cells were thawed and expanded in either DMEM (Gibco,11966025, [5 mM] glucose) or HPLM) supplemented with 5% HS (Gibco, 16050-122), 20 ng/mL epidermal growth factor (Peprotech), 10 µg/mL insulin (Sigma), 0.5 µg/mL hydrocortisone (Sigma), 100 ng/mL cholera toxin (Sigma, C8052-2MG), and 1x penicillin/streptomycin (Gibco). Subsequent experiments (proteomics, metabolomics, and lectin microarrays) were carried out in the same mediums with 5% dialyzed Fetal Bovine Serum (FBS) (Gibco, A3382001) replacing the standard HS used for MCF10A culture. Imaging experiments were carried out in both dialyzed FBS and HS with similar results obtained in either.

KHYG1 cells were provided by Alexandros Karampatzakis and David H. Raulet (UC Berkeley) and expanded in HPLM supplemented with 10% FBS, with recombinant human IL2 [40 U/mL] (Peprotech, 200-02). Co-cultures of KHYG1 and MCF10A cells were carried out in MCF10A medium, which did not affect the viability of KHYG1 cells within 24 h of transfer.

MB-MDA-231 tumor cells (ATCC) were grown in HPLM supplemented with 10% fetal bovine serum (FBS) (Hyclone) and 1x penicillin/streptomycin. HEK293T cells (ATCC) were maintained in DMEM supplemented with 10% FBS and 1x penicillin/streptomycin

HEK293T cells (ATCC) were maintained in DMEM supplemented with 10% FBS and 1x penicillin/streptomycin were used to produce lentiviral particles with psPAX2 (Addgene 12260), pMD2.G (Addgene 12259) and various transfer vectors described hereafter (https://www.addgene.org/guides/lentivirus/).

Human embryonic stem cell (hESC) line (H9, female) was provided by Susan Fisher (UCSF) and maintained in feeder-free conditions on tissue culture polystyrene coated with reconstituted basement membrane extract (rBM; Matrigel equivalent; R&D systems) suspended in DMEM (GenClone). hESCs were cultured in 50% Essential 8 (E8) media (Gibco) and 50% KnockOutTM Serum Replacement (KSR) media (E8/KSR) mix (Gibco) with 10 ng/ml bFGF (PeproTech) and 1x antibiotic-antimycotic (Gibco), referred to as hESCs growth media. Media was replaced every 24 h and cells were passaged every 3-4 days with 0.5 mM EDTA (Mankani) and 2% BSA (Fisher) in PBS, followed by resuspension in E8/KSR media supplemented with 10 μM Y27632 (ROCK inhibitor; TOCRIS). After 24 h post-passaging, media was switched to hESCs growth media without or without Y27632 on rBM coated glass coverslips for microscopy.

All cell lines were routinely tested and found to be free of mycoplasma contamination, and maintained in 5% CO_2_ at 37 °C. Experiments involving hESCs were approved by the University of California San Francisco Human Gamete, Embryo and Stem Cell Research Committee (UCSF GESCR) in the study numbered 11-05439.

### ROCK:ER

pBABEpuro3 ROCK:ER (*67*) was generously provided by Dr. Michael F. Olson, packaged into retroviral particles with phoenix cells, and used to generate stable MCF10A cells lines which were activated with 1 µg/mL 4-Hydroxytamoxifen (Sigma) in MCF10A culture medium.

### CMAS KO

Three non-overlapping gRNAs targeting a conserved, early exon were selected using Synthego’s CRISPR Design Tool and synthesized as modified sgRNAs (Synthego). The sgRNAs were resuspended to 100 μM in nuclease-free 10 mM Tris-EDTA pH 8.0, diluted to 100 μM in nuclease-free water, and complexed with 20 μM *S.py* Cas9 (Synthego) *in vitro* with a 9:1 molar ratio of sgRNA:Cas9 for 30 min prior to nucleofection. A suspension of approx. 100,000 MCF10a cells was nucleofected with program DS-138 using an Amaxa 4D-Nucleofector X using the SE Cell Line 4D X Kit S 32 RCT (Lonza V4XC-1032). Reactions were split between two 24-well plates and grown in complete media three days. Genomic DNA was extracted by first aspirating media from a 24-well plate of clones, treating each well with 100 μl of lysis buffer (0.45% NP40, 0.45% Tween20, 0.5x NEB Q5 PCR buffer, 0.1 mg/mL NEB Proteinase K) were incubated for 10 min at 55°C in and subsequently heated to 95°C for 10 min to inactivate the proteinase K. Then, 1 µL of DNA lysate was added to a 20 µL amplification reaction (Phusion Plus Green PCR Master Mix, ThermoFisher, F632L). 400-600 bp fragments flanking the edited regions were amplified, analyzed by gel electrophoresis, and sequenced (Elim Bio, Hayward, CA) to estimate edit efficiency using Inference of CRISPR Edits (ICE, Synthego). Then, from populations with est. >70% gene knock-out, clones were selected by limiting dilution in 96 well plates using MEGM Complete Medium (CC-3150, Lonza) with 100 ng/mL cholera toxin (Sigma, C8052). For each clone, gene knock-out was verified at the DNA level with sanger sequencing (Elim Bio, Hayward, CA) and at the protein level by immunoblot. Samples were run out on a 4-12% Bis-Tris SDS-PAGE Gel (BioRad, 3450125) at 180V for 65 min in 1x MOPS Buffer (BioRad, 1610788). Proteins were transferred onto 0.2 μm nitrocellulose membranes (BioRad, 1704271) using the Transblot Turbo System (BioRad, 1704150). Membranes were blocked in 1x Blocking Buffer (1x PBS pH 7.4, 0.05% Tween-20, 5% Bovine Serum Albumin) for 1h at room temperature with gentle agitation. Then, the membrane was washed thrice with PBS-T (1x PBS pH 7.4 + 0.05% Tween-20) allowing the membrane to shake for 5 min. Primary antibodies targeting the proteins were diluted in 1x Blocking Buffer to 1 µg/mL and incubated overnight at 4°C with gentle agitation. Following incubation in primary, the membranes were washed thrice with 1x PBS-T 0.05% for 5 min at room temperature. Then, a dilution of the appropriate secondary antibody (LiCor) was used to detect primary antibody binding. Finally, the blots were imaged with an Odyssey CLx Imager (LI-COR). Primary antibodies used: CMAS (Santa Cruz Biotechnology, (14W): sc-100486 and Thermo Fisher, PA5-89409), GNE (ThermoFisher, PA5-25238), NANS (ThermoFisher, PA553960), α-Tubulin (Thermo Fisher, T5168), Lamin A/C (Bethyl, A303-430A), GAPDH (Abcam, ab9485)

### ECM coated polyacrylamide hydrogel cell culture surfaces (PA-gels)

Cleaned (10% ClO, 1M HCL, then 100% EtOH) round #1 German glass coverslips (Electron Microscopy Services) were coated with 0.5% v/v (3-Aminopropyl)triethoxysilane (APTES, Sigma, 440140), 99.2% v/v ethanol, and 0.3% v/v glacial acetic acid for 2 h and then cleaned in 100% EtOH on an orbital shaker at 22 °C. APTES activated coverslips were coated with PBS buffered acrylamide / bis-acrylamide (Bio-Rad, 1610140 and 1610142) solutions (3% / 0.05% for 400 Pa, 7.5% / 0.07% for 6k Pa, and 10% / 0.5% for 60k Pa) polymerized with TEMED (0.1% v/v) (Bio-Rad, 1610801) and Potassium Persulfate (0.1% w/v) (Fisher, BP180) to yield a final thickness of ∼ 85 µm. PA-gels were washed with 70% EtOH and sterile PBS prior 3,4-dihydroxy-L-phenylalanine (DOPA, CAS 59-92-7, Alfa Aeser, A1131106) coating for 5 min at 22 °C protected from light with sterile filtered DOPA in pH 10 [10 mM] Tris buffer DOPA coated PA-gels were washed 2x with sterile PBS and ECM functionalized with 5 µg/mL human fibronectin (Millipore, FC010) in sterile PBS 1 h at 37 °C to generate an expected fibronectin coating density of 6 μM/cm^2^.

### Immunofluorescence microscopy

For lectin staining, cells/tissues were fixed in 4% paraformaldehyde (Electron Microscopy Services, 15710) in 1X PBS for 30 min at room temperature followed by subsequent PBS and Carbo-Free Blocking Solution (Vector, SP-5040-125) washes. Lectin staining was accomplished at [1:200] in Carbo-Free Blocking Solution at 4° overnight. Lectins/carbohydrate binding reagents: SNA-biotin (Vector, B-1305–2), GNL-biotin (Vector, B-1245-2), and WGA-rhodamine (Vector, RL-1022). SLBR-N-biotin was generated by expressing the glutathione S-transferase (GST)-NCTC10712BR fusion protein in *Escherichia coli* (strain BL21(DE3)) and purified using glutathione Sepharose Concentrated stock solutions [2 mg/mL] of SLBR-N were modified with Biotin via, EZ-Link NHS-Biotin (ThermoFisher, 20217), followed by thorough PBS dialysis at 4 °C, and then used as a staining reagent. For Siglec7-Fc staining, a precomplexing protocol [38] was used with 1 µg/mL Siglec7-Fc (R&D Systems, 1138SL050) was incubated with 1 µg/mL donkey anti-human-DyLight488 (Thermo-Fisher, PISA510126) in 1x Carbo-Free blocking solution and was incubated on an orbital shaker at 50 rpm for 15 min at RT prior to addition to sample.

For intracellular targets (e.g., cleaved caspase 3) cells/tissues were washed and blocked with a blocking buffer (HBSS fortified with: 10% FBS (Hyclone), 0.1% BSA (Fisher, BP1600), 0.05% saponin (EMD, L3771), and 0.1% Tween 20 (Fisher, BP337500). Primary antibodies [1:100-1:200] for 2 h at RT or 24 h at 4 °C, Secondary antibodies [1:1000] for 2 h at 22 °C. Antibodies used: Cleaved Caspase-3 (Asp175) (Cell Signaling, 9661, AB_2341188), CMAS (Sigma, HPA039905, AB_10795237), and HSF1 (Cell Signaling, 4356, AB_2120258).

Super resolution via optical reassignment (SoRa) confocal microscopy was accomplished with a Yokogawa CSU-W1/SoRa spinning disk confocal system in SoRa mode on a Ti2 inverted microscope stand (Nikon), and images acquired with an ORCA Fusion BT sCMOS camera (Hamamatsu).

Confocal microscopy was accomplished with a Nikon Eclipse Ti spinning disc microscope, Yokogawa CSU-X, Andor Zyla sCMOS, Andor Multi-Port Laser unit, and Molecular Devices MetaMorph imaging suite.

### Phasor FLIM

Phasor FLIM (*76*) was carried out on MCF10A cells (50-100 cells/mL) were seeded onto fibronectin coated PA-gels cast on 30 mm coverslips in 35 mm TC dishes. Mitochondria labeling was achieved by adding 2 µL of a tetramethyl rhodamine methyl ester (TMRM, ThermoFisher, T668) directly to the medium to achieve a final concentration of TMRM of 100 nM. The samples were then placed on the incubated and buffered stage (37 °C and 5% CO_2_) of a Zeiss LSM 880 confocal microscope equipped with a water immersion 40x 1.2 N.A. objective and MaiTai two photon laser (Spectra Physics). FLIM images were acquired by averaging 30 frames of 256×256 pixels at 16 us/pixel with a pixel size about 200-300 nm. The laser excitation was set at 740 nm with laser power optimized to have negligible photobleaching (<10% initial TMRM intensity). FLIM images were acquired using an A320 FastFLIM box device (ISS) coupled to Hamamatsu photo multipliers tube detectors (Hamamatsu, H7422P-40). The fluorescence signal was acquired through a 690 nm dichroic filter and directed to the detectors. NADH signal was acquired through a band pass filter (460/80 nm) and TMRM signal was acquired through a long pass filter (520 nm) and band pass filter (641/75 nm). The signal was processed by SIM FCS software (SIM FCS 4, https://biii.eu/globals-images-simfcs-4) developed by the Laboratory for Fluorescence Dynamics, University of California, Irvine. The Phasor transformation was carried out using SIM FCS software calibrating the instrument response with a 100 nM Coumarin 6 solution (mono exponential lifetime at 2.5 ns) as a reference for FLIM. A systematic and automated methodology was used to process and analyze fluorescence lifetime images by way of a custom Python script (https://github.com/aelefebv/lfdfiles). The workflow comprises several steps: file conversion, image segmentation, masking, G (real) and S (imaginary) components calculation, bound NADH fraction determination, and results exportation as CSV files. Initially, raw R64 files were converted to TIFF format for compatibility and image manipulation. The TMRM channel image underwent segmentation to distinguish mitochondrial from cytosolic signals, using a diffuse background subtraction removal technique, as described in our previous paper (*77*). Masks were applied to corresponding phase and modulation images, enabling the extraction of cellular compartment-specific FLIM information. G and S parameters, key fluorescence lifetime indicators, were calculated. Bound fluorophore fractions (FB) were calculated, providing insight into NADH metabolic trajectory within cellular compartments. Mean FB values were calculated for each sample and exported as CSV files. Additionally, the script generated a composite colormap image visualizing NADH FB values and exported G and S coordinates as CSV files for downstream analysis. This custom Python script facilitated comprehensive processing and analysis of fluorescence lifetime images, ensuring reproducibility and accuracy in examining cellular interactions and microenvironments.

### Proteomics (timsTOF)

1 million cells were seeded on round 50 mm^2^ varied stiffness ECM coated PA-gels cultured for 24 h in HPLM or DMEM based MCF10A medium. Cells were washed with sterile PBS once, then cells were detached with cold PBS and a cell scraper (rubber policeman). Samples were prepared using the Sample Preparation by Easy Extraction and Digestion (SPEED) method. (*78*). Briefly, cells were lysed through the addition 1:4 (v/v) of > 99% trifluoruacetic acid (TFA) for 3 minutes. Neutralization buffer (2M TrisBase) was added at 10x the volume of added TFA, followed by the addition of reduction/alkylation buffer (100 mM Tris(2-carboxyethyl)phosphine (TCEP), 400 mM 2-Chloroacetamide (CAA) in H2O) at 1.1x the volume of added TFA. Samples were incubated at at 95 °C for 5 min. Protein concentration was determined using Bradford assay (Thermo) according to the manufacturer’s directions. Trypsin (Promega) was added at a 1:50 enyme:protein ratio and digested for 20 hours at 37 °C and 1000 rpm.The samples were then acidified with Trifluoroacetic acid (TFA) [∼1% final] to yield a sample pH of 2. Samples were then desalted on Ultra Micro C18 Spin Column (Nest Group). Samples were resuspended in 0.1% formic acid (FA) and analyzed by DDA-PASEF on a timsTOFpro (Bruker) (*79*) interfaced with an EASY-nLC 1000 (Thermo) Samples were separated on a 25 cm pepSep column over a 55 minute gradient at 0.5 µL/min. Following loading, peptides were analyzed over 65 minutes at a gradient of 5 to 30% B (0.1% FA in 80% acetonitrile (ACN), followed by a 1 minute ramp to 90% B at high flow (1 µL/min). B was held a 90% for 4 minutes, then ramped down to 5%. Peptides were sprayed through a 20 mm ZDV emitter held at 1700 V and 200°C. Samples were analyzed in positive ion mode using DDA-PASEF over a 400-1550 m/z range. Raw files were analyzed using the Andromeda search engine (*80*) in MaxQuant using the default TIMS-DDA settings (*81*). Variable modifications were set to oxidation of methionine and N-terminal acetylation, with carbamidomethylation of cysteines set as a fixed modification. Up to two missed tryptic cleavages were permitted. Peptide and protein identifications were filtered to a 1% false-discovery rate (*82*). Searches were performed against a protein database of the human proteome (downloaded from Uniprot on 02/03/20). Quality control analysis was performed via artMS (http://artms.org) and statistical testing was performed with MSstats (*83*).

### LC-MS Metabolomics and ^13^C6-Glucose LC-MS Metabolomics

1 million cells were seeded on 50 mm^2^ varied stiffness ECM coated PA-gels cultured for 22 h in 5 mM glucose DMEM of HPLM based MCF10A media. The media was exchanged for media with or without 5 mM ^13^C6-Glucose (Cambridge Isotope Laboratories Inc., CLM-1396) in HPLM or DMEM based MCF10A media for 2 h (for ^13^C6-Glucose media, no-glucose DMEM or bespoke no glucose HPLM was used as a base medium that lacked ^12^C-Glucose). Cells were washed twice with PBS and extracted with mass spectrometry grade 80% methanol (ThermoFisher, A456-1) and 20% water (ThermoFisher, W6500) supplemented with 1 nmol DL-Norvaline (Sigma, N7502). Protein concentrations of the methanol extract was determined via BCA (Pierce, 23225) with no significant variability assessed (5 uL transferred into 45 uL RIPA buffer, 5 uL of the RIPA dissolved solution assayed). Insoluble material was pelleted in a 4°C centrifuge at 16k x g, supernatant was transferred and dried in a Speedvac. Dried metabolites were resuspended in 50% ACN:water and 1/10th of the volume was loaded onto a Luna 3 um NH2 100A (150 × 2.0 mm) column (Phenomenex). The chromatographic separation was performed on a Vanquish Flex (Thermo Scientific) with mobile phases A (5 mM NH_4_AcO pH 9.9) and B (ACN) and a flow rate of 200 μL/min. A linear gradient from 15% A to 95% A over 18 min was followed by 9 min isocratic flow at 95% A and reequilibration to 15% A. Metabolites were detected with a Thermo Scientific Q Exactive mass spectrometer run with polarity switching (+3.5 kV / -3.5 kV) in full scan mode with an m/z range of 65-975. TraceFinder 4.1 (Thermo Scientific) was used to quantify the targeted metabolites by area under the curve using expected retention time and accurate mass measurements (< 5 ppm). Values were normalized to cell number and sample protein concentration. Relative amounts of metabolites were calculated by summing up the values for all isotopologues of a given metabolite. Fractional contributional (FC) of ^13^C carbons to total carbon for each metabolite was calculated (*84*). Data analysis was accomplished using in-house developed R scripts.

### Metaboverse

Reaction representations and multi-condition reaction component comparisons were prepared using Metaboverse (v0.9.0) (*30*). Input proteomics and metabolomics were prepared by calculating the log_2_ fold change and Benjamini-Hochberg corrected Student’s T-test statistic for each comparison.

### Scanning angle interference microscopy (SAIM)

Silicon wafers with ∼1900 nm thermal oxide (Addison Engineering) were diced into 7 mm by 7 mm chips, and the oxide layer thickness for each chip was measured with a FilMetrics F50-EXR. Prior to use, the chips were then cleaned with 1:1 methanol and hydrochloric acid for 15 minutes, followed by 1 minute of plasma cleaning (Harrick Plasma, PDC-001). The chips were incubated with 4% (v/v) (3-mercaptopropyl)trimethoxysilane in 100% ethanol for 30 minutes at RT. After washing with 100% ethanol, the chips were subsequently incubated with 4 mM 4-maleimidobutyric acid N-hydroxysuccinimide ester in 100% ethanol for 30 minutes, then rinsed with PBS. 50 µg/ml human plasma fibronectin (AF-647 conjugated) in PBS was incubated with the functionalized chips overnight at 4 °C. Cells were seeded onto the chips at 2 × 10^4^ cells/cm^2^ in full culture medium and incubated for 24 hours. Samples were then rinsed 3x in PBS, inverted into a 35 mm glass-bottom imaging dish containing imaging buffer and imaged at 37 °C. SAIM was conducted on a custom circle-scanning microscope (*85*). The core of the setup was an inverted fluorescence microscope (Ti-E, Nikon). The excitation lasers (488 nm, Coherent; 560 nm, MPB Communications Inc.; and 642 nm, MPB Communications Inc.) were combined into a colinear beam by a series of dichroic mirrors (Chroma). The combined output beam was attenuated and shuttered by an AOTF (AA Opto-Electronic). The beam was directed onto a pair of galvanometer scanning mirrors (Cambridge Technology). The image of the laser on the scanning mirrors was magnified and relayed to the sample by two 4 f lens systems, a beam expanding telescope and a scan lens / objective lens combination. The beam expander was formed by f = 30 mm and f = 300 mm achromatic lenses with a m = 1 zero-order vortex half-wave plate positioned between them and positioned 2 f from the 300 mm achromatic scan lens (Thorlabs). SAIM experiments were performed with a 60x 1.27 N.A. water immersion objective (Nikon). Fluorescence emission was collected with a quad band filter cube and single band filters (TRF89901-EMv2, Chroma) mounted in a motorized filter wheel (Sutter). Images were acquired with a Zyla 4.2 sCMOS (Andor) camera or an iXon 888 EMCCD (Andor) using the microscope’s 1.5x magnifier for a total magnification of 90X. The open-source software Micro-Manager was used for camera and filter wheel control and image acquisition. The circle-scanning galvometers were operated in an autonomous fashion using a custom-designed 16-bit PIC microcontroller (*85*). 32 images were acquired at varying incidence angles for the circle-scanned excitation beam. During a typical image acquisition sequence, changes in the scanned incidence angle were triggered by a TTL signal from the camera to the microcontroller. The incidence angles were evenly spaced from 5 to 43.75 degrees with respect to the wafer normal in the imaging media. To obtain the reconstructed height topography of the samples, the raw image intensities, I_j_, at each incidence angle, θ_j_, were fit pixelwise by nonlinear least-squares to an optical model:

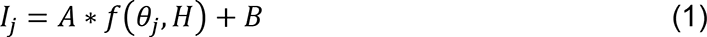

where H is the unknown sample height and A and B are additional fit parameters. The vortex half-wave plate in the optical setup maintained the s-polarization of the circle-scanned excitation laser. For s-polarized monochromatic excitation of wavelength, λ, the probability of excitation, *f*(*θ*_j_, *H*), for the system is given by:

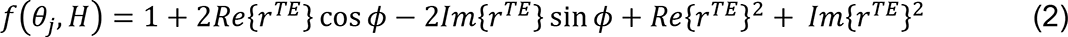

where the phase shift, ϕ, and the reflection coefficient for the transverse electric wave, *r*^TE^, are given by:

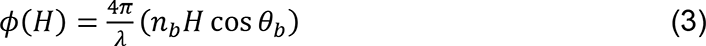

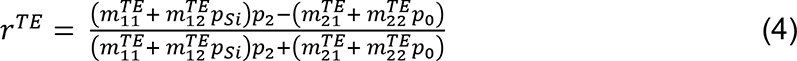

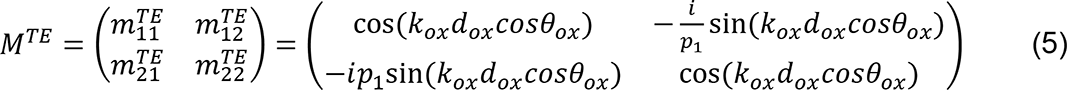

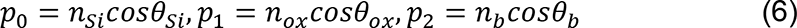

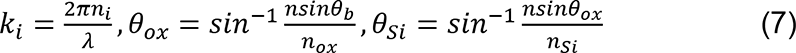

where *k*_i_ is the wavenumber in material i; *n*_Si_, *n*_ox_ and *n*_b_ are the refractive index of the silicon, silicon oxide and sample, respectively; *θ*_Si_, *θ*_ox_ and *θ*_b_ are the angles of incidence in the silicon, silicon oxide and sample, respectively; and *d_ox_* is the thickness of the silicon oxide layer. The angles of incidence in silicon oxide and silicon were calculated according to Snell’s Law. To quantify the glycocalyx thickness of cells, the average height above the silicon substrate was calculated for a 100 x 100 pixel subregion in each cell. The glycocalyx thickness was reported as the height of the MemGlow dye 560 signal minus the height of the AF647-fibronectin layer on the silicon substrate. Packages for fitting SAIM image sequences with the above model have been implemented in C++ and Julia and are available through GitHub: (https://github.com/mjc449/SAIMscannerV3 and https://github.com/paszeklab/SAIMFitKit.jl)

### Parallel Plate Shear Rheology

Viscoelastic properties of tissues were determined using an oscillatory rheometer (Anton-Paar, M302e) with parallel-plate geometry (8 mm) and a gap height of ∼ 0.2 mm under 0.1% strain and 1 Hz frequency at 37 °C in a humidity-controlled chamber.

### _HPLM_DMEM Salt

HPLM was generated in-house (*24*) with the inorganic salt composition of DMEM (CaCl_2_ [1.802 mM], Fe(NO_3_)_3_, [0.248 mM]; KCl, [5.333 mM]; MgSO_4_, [0.814 mM]; NaCl [110.345 mM]; NaHCO_3_, [44.048 mM], NaH_2_PO_4_ [0.906 mM]). Cells were cultured for at least 7 days in HPLM^DMEM Salt^ prior to comparisons to cells in HPLM. We did not find that MCF10A cells cultured in HPLM^DMEM Salt^ had proliferative differences to HPLM; however, they did spread more at low confluence than HPLM on tissue culture polystyrene surfaces.

### Flow Cytometry (WGA staining)

Cells were trypsinized, washed once with cold FACS buffer (PBS with 10mM HEPES, 5% BSA), and stained on ice with Rhodamine labeled WGA (1:500) for 20 minutes. After washing, cells were analyzed on BD LSR II flow cytometer, collecting approximately 5000 events per sample. Data were analyzed with FlowJo (v10) to quantify mean fluorescence intensity of WGA staining on live single cells.

### Osmolality

Osmolality of culture medium was determined by freezing-point depression with an Osmo1 analyzer (Advanced Instruments).

### Hsf1 Conditional KO Mouse Model

*Hsf1^flox^* mice (*57*) were generously provided by Chengkai Dai, NIH. *Hsf1*^flox^ mice were crossed with MMTV-Cre/Line D (JAX labs, 003553) and backcrossed to stain purity with with C57BL/6J mice (JAX labs, 000664). Floxed alleles were assessed with genotyping primers: P_1_ (5′GGGTATGGGGGACTTTTAGG-3′), P_2_ (5′-AGTGAGGCCCATGTAACCAG-3′), P_3_ (5′-TCCTCCTCCCTCCCAAGTGGG-3′).

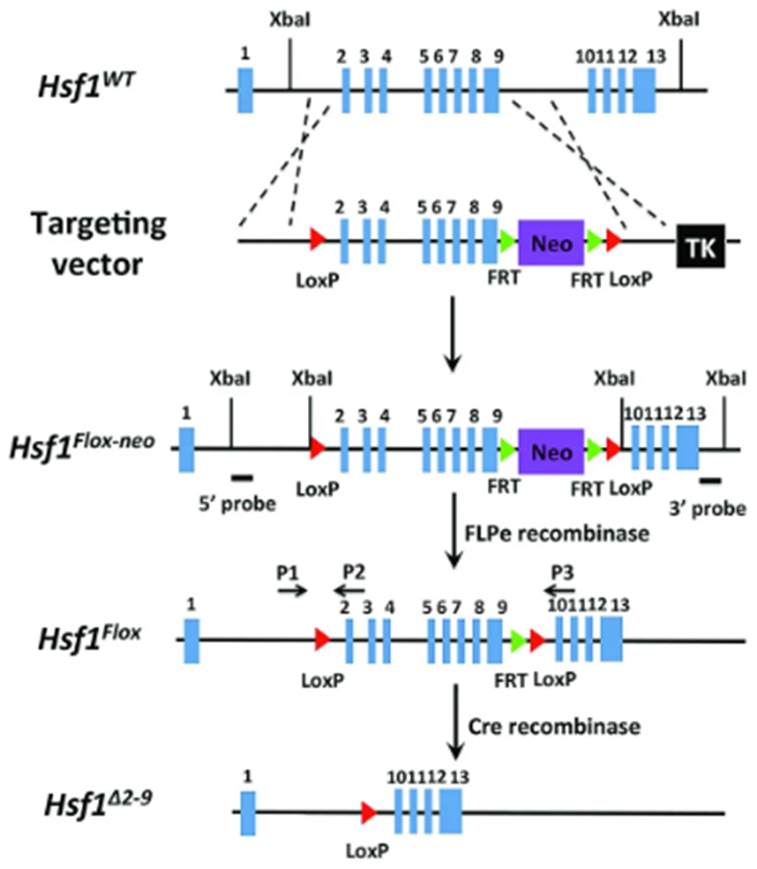

### Lectin Microarray

Flash frozen MCF10A cell pellets were washed with protease inhibitor cocktail supplemented PBS and sonicated on ice until homogenous. 20 μg of protein from the homogenized samples were then labeled with Alexa Fluor 555-NHS (*86*). A reference sample was prepared by pooling equal amounts (by total protein) of all samples and labeling with Alexa Fluor 647-NHS (Thermo Fisher). Table S1 summarizes the print list for the experiment Printing, hybridization, and data analysis (*87*).

**Table S1.**
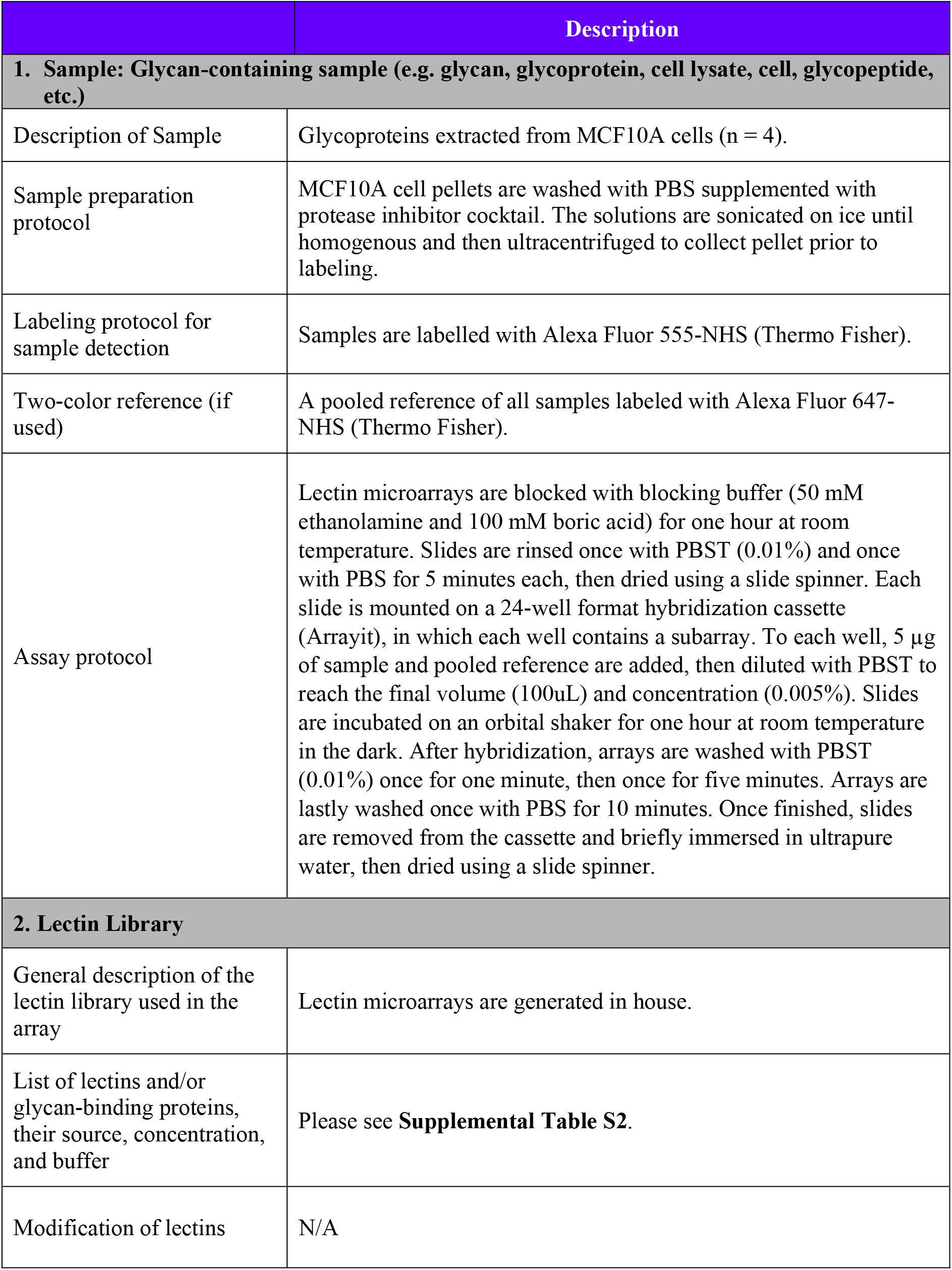
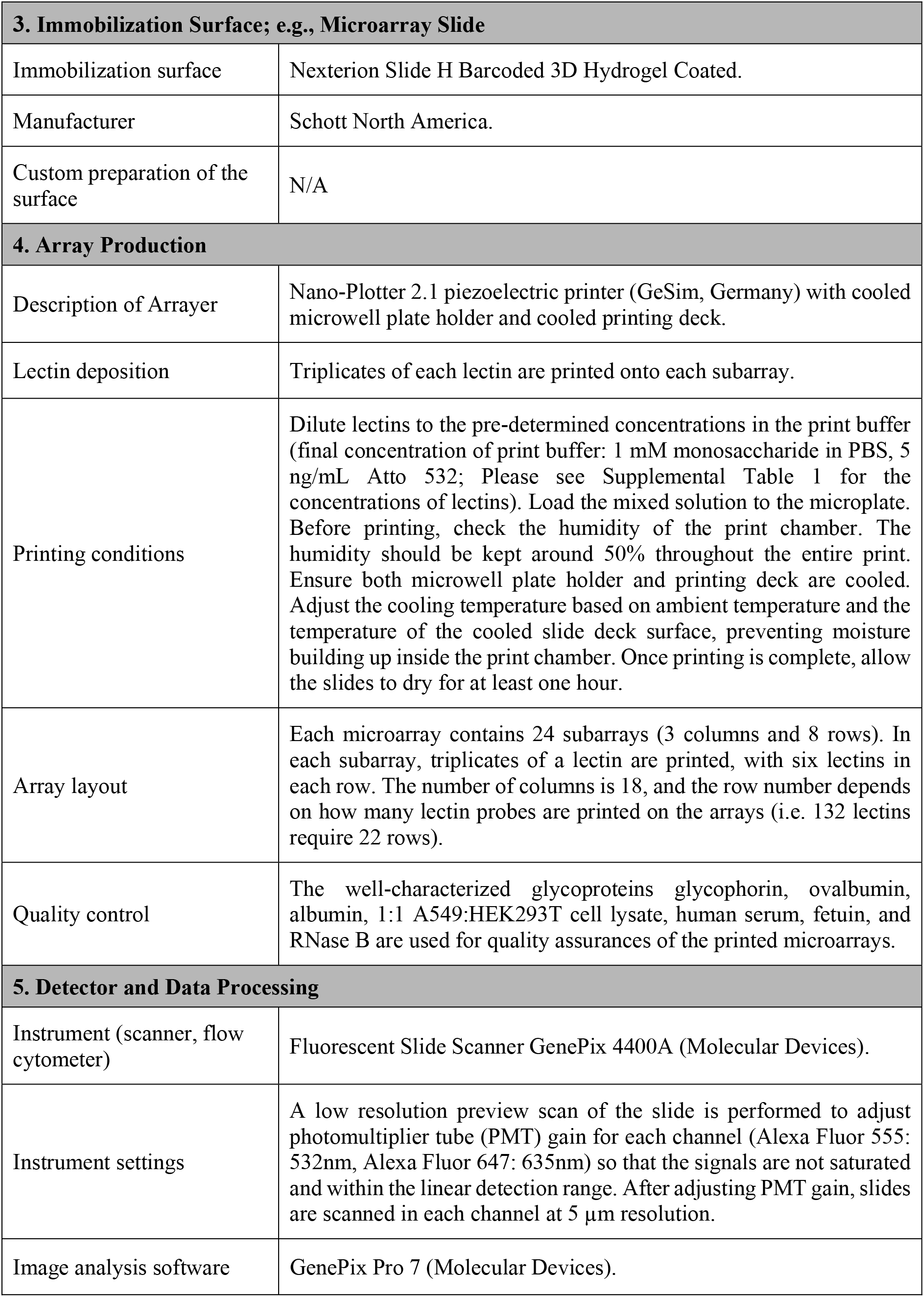
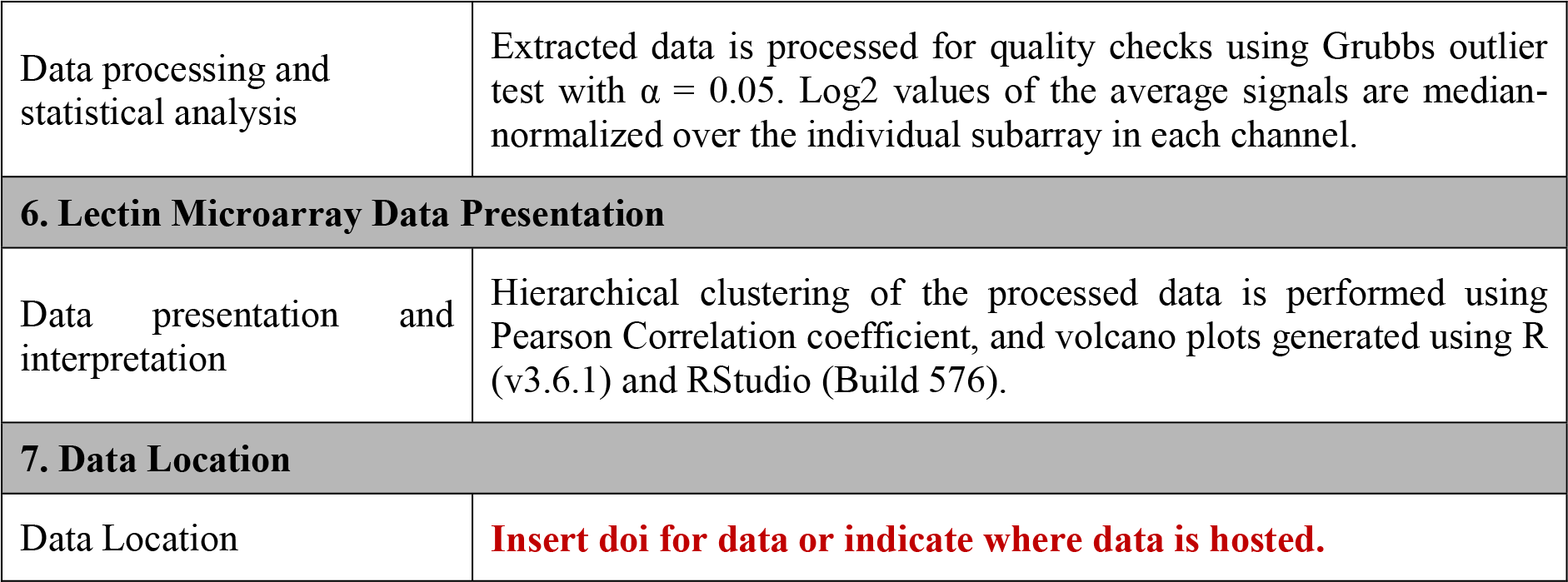
Lectin Microarray Information.

**Table S2.** Lectins used in microarrays.

| Lectin | Species/Origin | Print Conc. ($\mu\text{g/mL}$) | Rough Specificity /Inhibitory monosaccharide | Vendor/Source |
| --- | --- | --- | --- | --- |
| AAL | *Aleuria aurantia* | 2000 | Fucose | Vector |
| ACA | *Amaranthus Caudatus* | 2000 | Gal-β1,3-GalNAc | EY |
| AIA | *Artocarpus integrifolia* | 2000 | β1,3-GalNAc | Vector/EY/Glycomatrix |
| AMA | *Allium moly* | 2000 | Oligo mannose | EY |
| Anti-B.G.A | MAB mouse IgM [HE-193] | undiluted | Blood group A antigen | Thermo Fisher |
| Anti-B.G.B | MAB mouse IgM [HEB-29] | undiluted | Blood group B antigen | Abcam/Thermo Fisher |
| Anti-E-selectin | MAB mouse IgG1 [P2H3] | undiluted | E-selectin | Thermo Fisher |
| Anti-Ficolin-1 | MAB mouse | undiluted | Ficolin-1 antigen | Lifespan Biosciences |
| Anti-Ficolin-3 | MAB mouse IgG1 [4H5] | undiluted | Ficolin-3 antigen | Lifespan Biosciences |
| Anti-Galectin-3 | MAB mouse IgG1 [A3A12] | undiluted | Galectin-3 antigen | Thermo Fisher |
| Anti-B.G.H1 | MAB mouse IgG3 [17-206] | undiluted | Blood group H1 antigen | Thermo Fisher |
| Anti-B.G.H2 | MAB mouse IgM [A46-B/B10] | undiluted | Blood group H2 antigen | Santa Cruz Biotechnology |
| Anti-IgM | MAB mouse IgG1 [SA-DA4] | undiluted | IgM | Thermo Fisher |
| Anti-B.G. Lewis A | MAB mouse IgG1 [7LE] | undiluted | Lewis A | Abcam/Thermo Fisher |
| Anti-B.G. Lewis B | MAB mouse IgM [2-25LE] | undiluted | Lewis B | Abcam/Sigma |
| Anti-B.G. Lewis Y | MAB mouse IgM [F3] | undiluted | Lewis Y | Abcam |
| Anti-Lewis X [P12] | MAB mouse IgM [P12] | undiluted | Lewis X | Sigma |
| Anti-Mac-2bp | MAB mouse IgG1 [SP2] | undiluted | LGALS3BP | Thermo Fisher |
| Anti-MBL | MAB mouse IgG1 [3B6] | undiluted | Collectin-1 | Abcam |
| Anti-P-Selectin | MAB mouse IgG1 [Psel.KO2.3] | undiluted | P-selectin | Thermo Fisher |
| AOL | *Aspergillus oryzae* | 2000 | Fucose | TCI America |
| ASA | *Allium sativum* | 2000 | Mannose | EY |
| BambL | *Burkholderia ambifaria* | 2000 | Fucose | Generated in house |
| BC2L-A | *Burkholderia cenocepacia* | 2000 | Mannose | Elicityl |
| Blackbean | *Blackbean crude* | 2000 | GalNAc | EY |
| BPA | *Bauhinia purpurea* | 2000 | β-Gal / β-GalNAc | Vector |
| CA | *Colchicum autumnale* | 2000 | Bi-antennary N-linked glycans | EY |
| ConA | *Canavalia ensiformis* | 2000 | Tri-mannose core | Vector/Thermo Fisher |
| CSA | *Cystisus scoparius* | 2000 | Terminal GalNAc | EY |
| DBA | *Dolichos biflorus* | 2000 | GalNAc | Vector |
| diCBM40 | engineered NanI from *Clostridium perfringens* | 1500 | $\alpha$ Sialylation | Generated in house |
| DSA | *Datura stramonium* | 2000 | LacNAc | Vector/Sigma |
| ECA | *Erythrina cristagalli* | 2000 | LacNAc | Vector |
| GNA/GNL | *Galanthus nivalis* | 2000 | Oligo mannose | Vector/Sigma |
| GS-I | *Griffonia simplicifolia-I* | 2000 | α-Gal / Lac | Vector |
| GS-II | *Griffonia simplicifolia-II* | 2000 | GlcNAc | Vector |
| H84T | *Banana lectin* | 1000 | High mannose | Gift from Dr. David Markovitz |
| HHL | *Hippeastrum Hybrid* | 2000 | Oligo/High mannose | Vector/BioWorld |
| HPA | *Helix pomatia* | 2000 | Blood Group A | Sigma |
| LcH | *Lens culinaris* | 2000 | Core Fucose | Vector/Aniara Diagnostica/EY/Medicago |
| LEA/LEL | *Lycopersicon esculentum* | 2000 | GlcNAc | Vector |
| Lotus | *Lotus tetragonolobus* | 2000 | Fucose | Vector/EY |
| MAL-I | *Maackia amurensis-I* | 2000 | Sialylation/Sulfation | Vector |
| MAL-II | *Maackia amurensis-II* | 2000 | Sialylation/Sulfation | Vector |
| MNA-G | *Morus nigra Morniga G* | 2000 | GalNAc | EY |
| MNA-M | *Morus nigra Morniga M* | 2000 | Oligo mannose / Gal | EY |
| MPA/MPL | *Maclura pomifera* | 2000 | β1,3-GalNAc | Vector |
| NPA | *Narcissus pseudonarcissus* | 2000 | Oligo mannose | Vector |
| PHA-E | *Phaseolus vulgaris Erythroagglutinin* | 2000 | Bisecting GlcNAc | Vector/EY |
| PHA-L | *Phaseolus vulgaris Leukoagglutinin* | 2000 | β1,6 Branching N-Link glycans | Vector/EY |
| PNA | *Arachis hyogaea* | 2000 | Gal-β1,3-GalNAc | Vector |
| Recombinant Protein A | *Staphylococcus aureus* | 2000 | Immunoglobulins | Thermo Fisher |
| Recombinant Protein G | *Streptococcus* Group G | 2000 | Immunoglobulins | Thermo Fisher |
| Recombinant Protein L | *Peptostreptococcus magnus* | 2000 | Immunoglobulins | Thermo Fisher |
| PSA | *Pisum sativum* | 2000 | Core Fucose | Vector/Glycomatrix |
| PSL | *Polyporus squamosus* | 2000 | α2,6 sialylation | EY |
| PTL-II | *Psophocarpus tetragonolobus-II* | 2000 | α2 Fucose | Vector |
| RCA120 | *Ricinus Communis Agglutinin I* | 2000 | Gal / Lac | Vector |
| rGRFT | *recombinant Griffithsin* | 1700 | High mannose | Gift from Dr. Barry O'Keefe |
| Ricin B Chain | *Ricinus communis* | 2000 | Gal | Vector |
| SBA | *Glycine max* | 2000 | LacdiNAc | Vector |
| SLBR-B | *Streptococcus gordonii M99* | 2300 | α2,6 sialylation | Generated in house |
| SLBR-H | *Streptococcus gordonii DL1* | 2000 | α2,3 sialylation | Generated in house |
| SLBR-N | *Streptococcus gordonii UB10712* | 2000 | α2,3 sialylation | Generated in house |
| SNA | *Sambucus nigra* | 2000 | α2,6 sialylation | Vector |
| SNA-II | *Sambucus nigra-II* | 2000 | α2 Fucose /oligo mannose | EY |
| TJA-II | *Trichosanthes japonica-II* | 2000 | α2 Fucose | Aniara Diagnostica |
| TL | *Tulipa sp.* | 2000 | GlcNAc | EY |
| UDA | *Urtica dioica* | 2000 | GlcNAc / Oligo mannose | EY |
| UEA-I | *Ulex europaeus-I* | 2000 | α2 Fucose | Vector/Sigma |
| UEA-II | *Ulex europaeus-II* | 2000 | GlcNAc | EY |
| VVA | *Vicia villosa* | 2000 | Terminal GalNAc | Vector |
| WFA | *Wisteria floribunda* | 2000 | GalNAc-β1,4 | Vector |
| WGA | *Triticum vulgare* | 2000 | GlcNAc | Vector |

## Supplemental Figures

**Figure S1:**
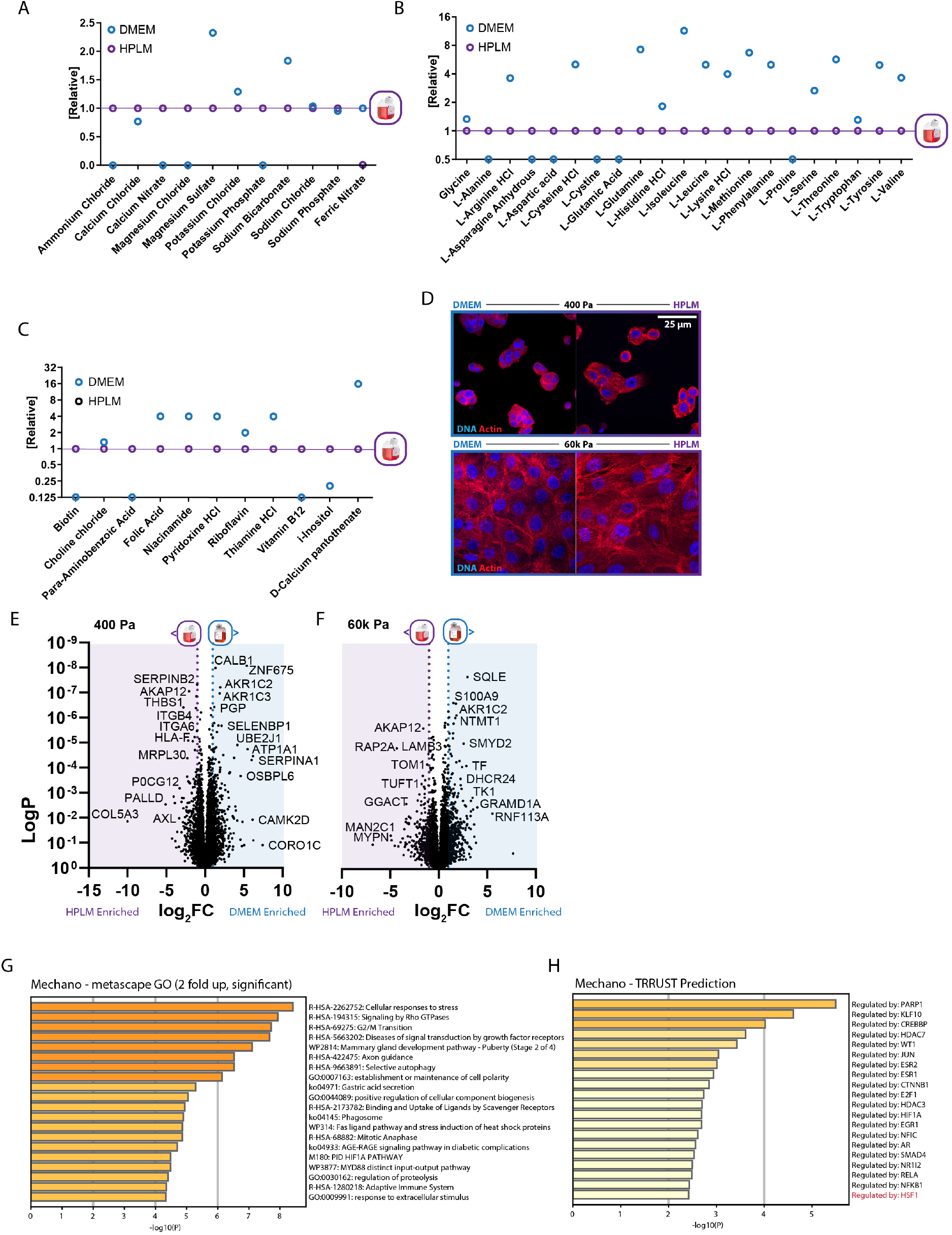
A. Relative abundance of salts in HPLM or DMEM. B. Relative abundance of amino acids in HPLM or DMEM. C. Relative abundance of vitamins and cofactors in HPLM or DMEM. D. Representative confocal microscopy of phalloidin (red) and DAPI (blue) staining of MCF10A cells on 400 Pa or 60k in DMEM of HPLM. E. Volcano plot depicting relative abundance of proteins in MCF10A cells cultured on 400 Pa in HPLM vs DMEM (fold change, DMEM/HPLM). F. Volcano plot depicting relative abundance of proteins in MCF10A cells cultured on 60k Pa in HPLM vs DMEM (fold change, DMEM/HPLM). G. Top GO categories representative of the significantly induced (two fold and up) proteins in MCF10A cells cultured on 60k vs 400 Pa in HPLM. H. TRRUST-based prediction of which transcription factors facilitate expression of the proteins significantly induced (two fold and up) in MCF10A cells cultured on 60k vs 400 Pa in HPLM.

**Figure S2:**
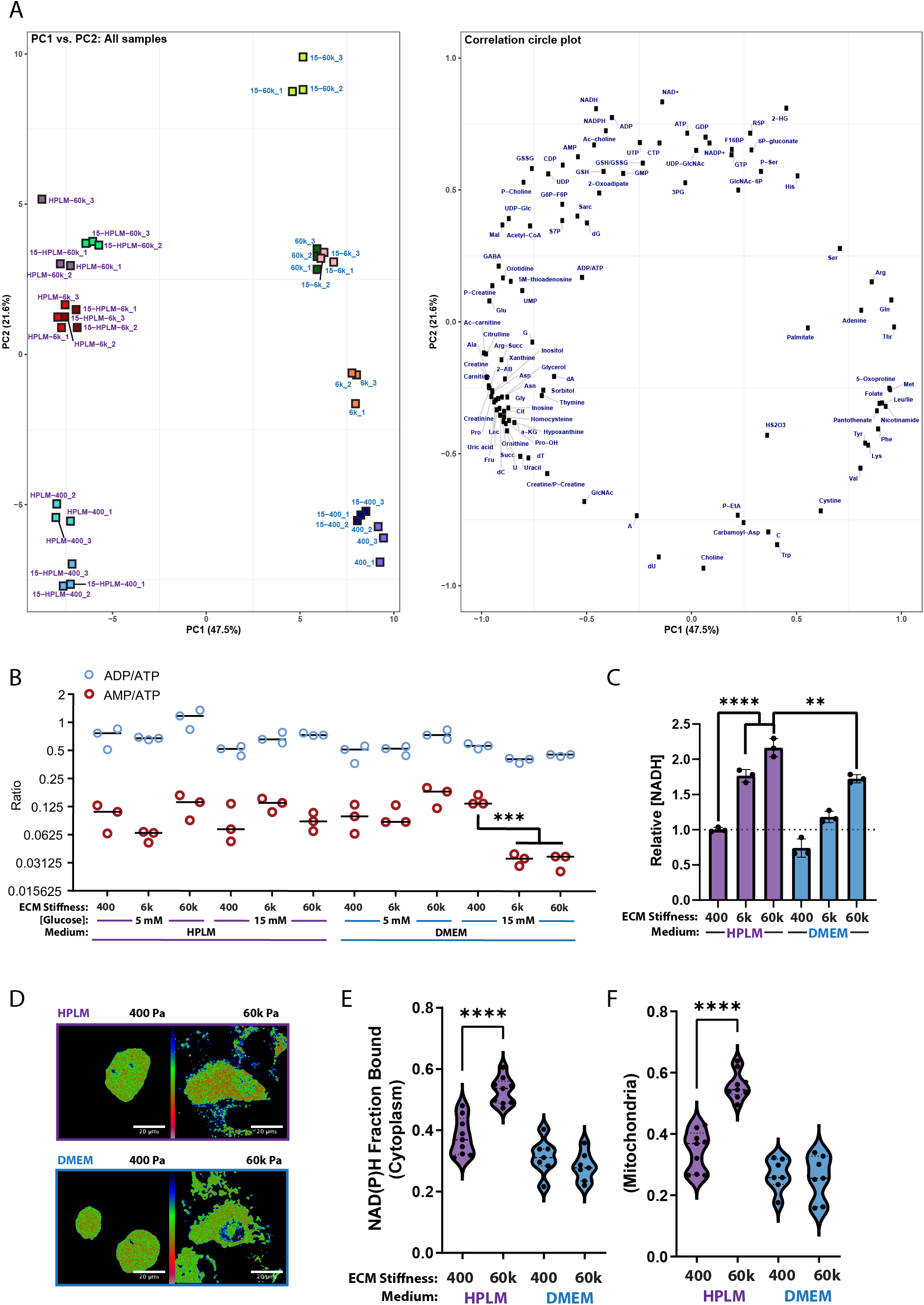
A. Principal Component Analysis (PCA) of intracellular metabolite levels of MCF10A cells cultured on 400, 6k, and 60k Pa in HPLM or DMEM with 5 or 15 mM glucose (indicated as 15- on plot), metabolites contributing to PC1 or 2 plotted on the right (n=3 biological replicates, associated with Fig. 2A-B) B. AMP/ATP and ADP/ATP concentration ratios between MCF10A cells cultured on 400 Pa, 6k Pa, or 60k Pa in HPLM or DMEM with 5 or 15 mM glucose. (n=3 biological replicates, associated with Fig. 2A-B) C. Relative abundance of NADH between MCF10A cells cultured on 400 Pa, 6k Pa, or 60k Pa in HPLM or DMEM with 5 or 15 mM glucose. (n = 3 biological replicates, associated with Fig. 2A-B) D. Representative images of the fraction of bound NADH, calculated using the phasor approach to the fluorescence lifetime imaging (FLIM) analysis of NADH, of MCF10A cells in HPLM (top) and DMEM (bottom) media, and on 400 Pa (left) and 60 kPa (right) conditions. E. The mean cytoplasmic fraction of bound NADH, where each point represents the average fraction bound in one field of view. (n = 7, 9 fields of view for DMEM, HPLM conditions, respectively) F. The mean mitochondrial fraction of bound NADH, where each point represents the average fraction bound in one field of view. (n = 7, 9 fields of view for DMEM, HPLM conditions, respectively)

**Figure S3:**
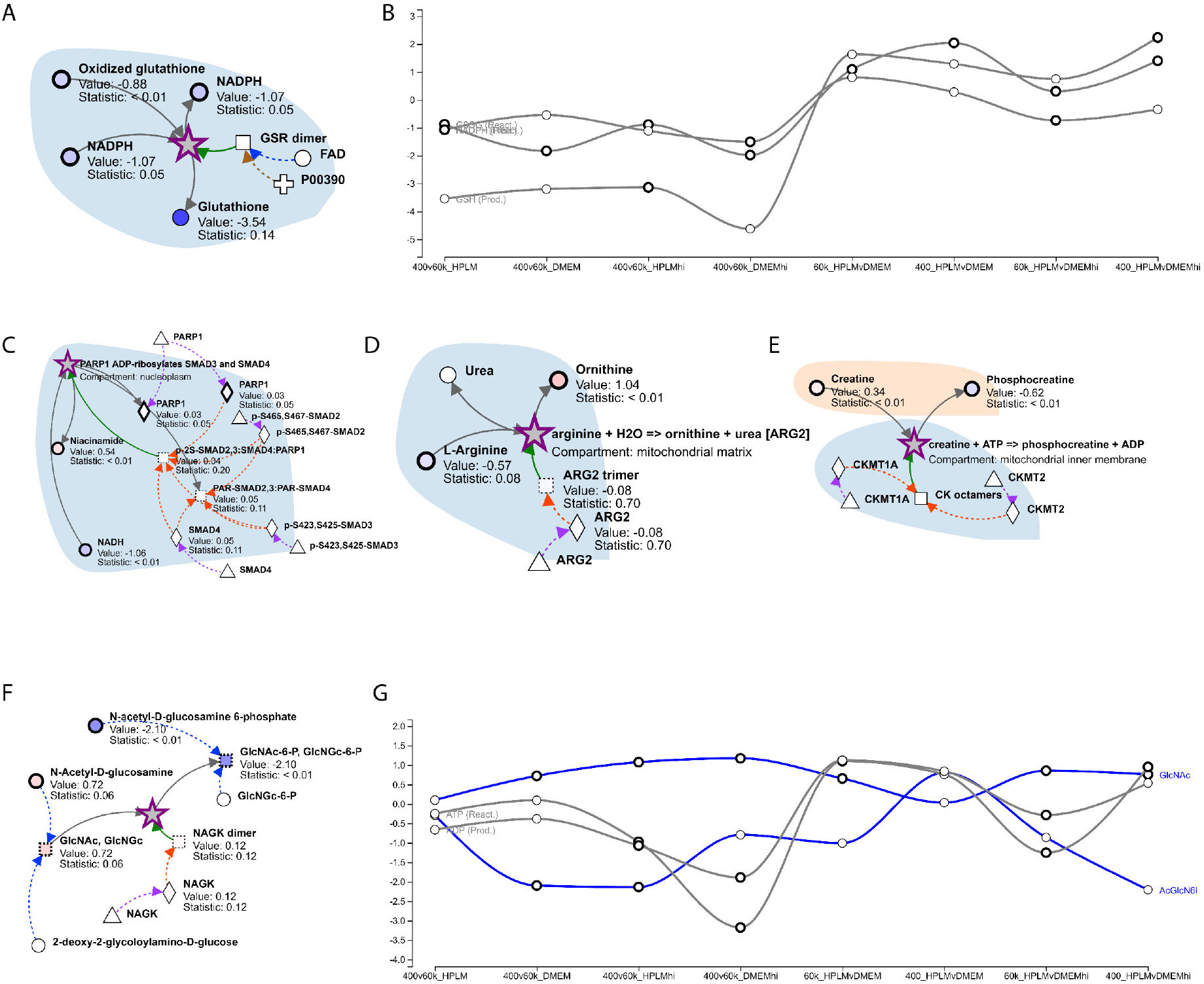
A. Predicted reaction pattern to be enriched as a product of 60k Pa over 400 Pa microenvironments in HPLM more strongly than in DMEM. B. Multiple comparison line plot of the relative abundance of products and reactants contributing to the prediction in A. Hi indicates 15 mM glucose. C. (C-E) Predicted reaction pattern to be enriched as a product of 60k Pa over 400 Pa microenvironments, regardless of medium. G. Predicted reaction pattern to be enriched as a product of 60k Pa over 400 Pa microenvironments in DMEM more strongly than in HPLM. H. Multiple comparison line plot of the relative abundance of products and reactants contributing to the prediction in F. Hi indicates 15 mM glucose.

**Figure S4:**
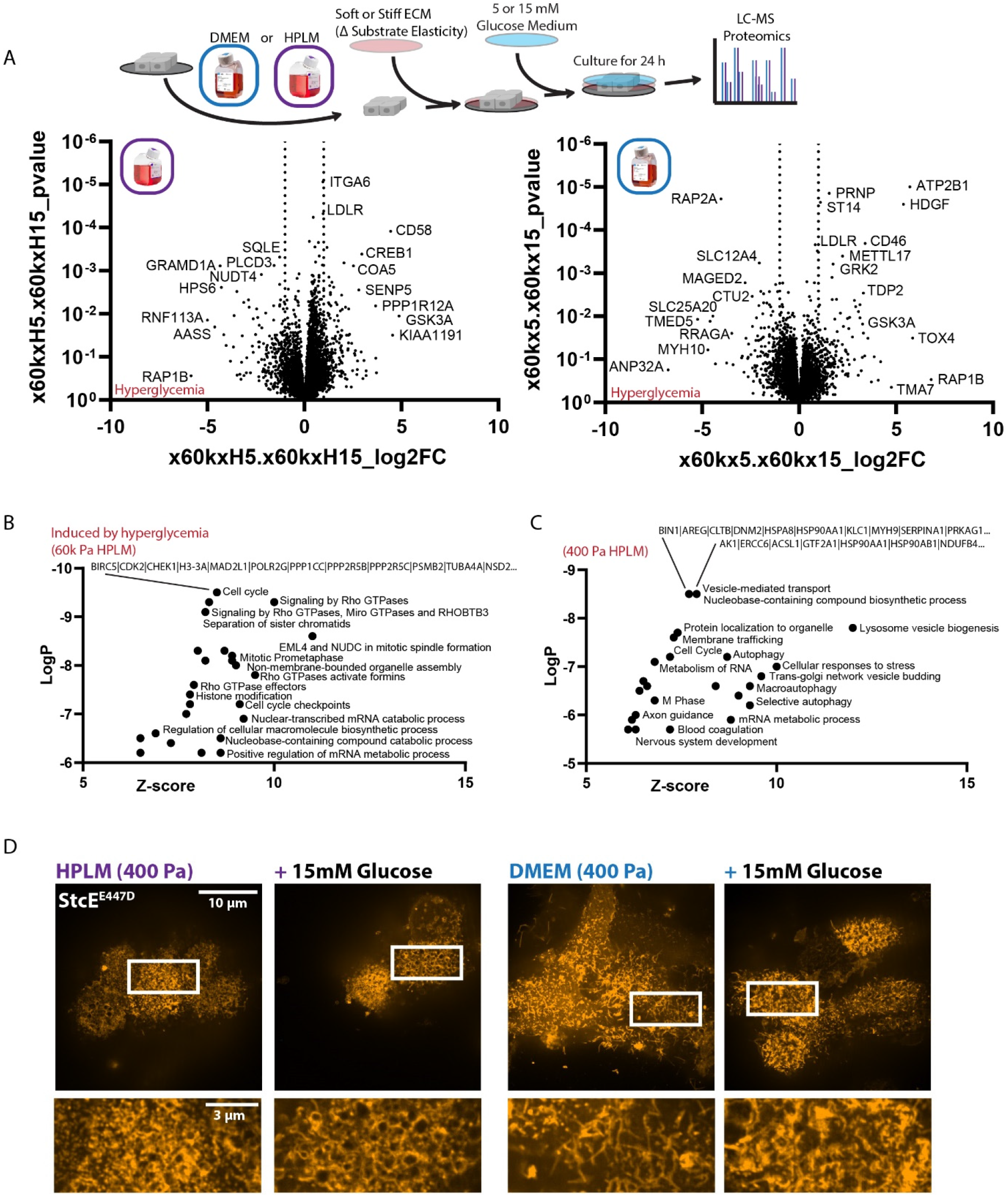
A. Graphical representation of experimental design related to A-C and volcano plots of fold change abundance of proteins in response to hyperglycemia (enriched to left) on 60k Pa in a given medium (HPLM, left plot, purple and DMEM, right plot, blue). (n = 3 biological replicates) B. Top GO categories representative of the significantly induced proteins in MCF10A cells cultured in hyperglycemia on 60k Pa in HPLM C. Top GO categories representative of the significantly induced proteins in MCF10A cells cultured in hyperglycemia on 400 in HPLM. D. Representative SoRa confocal microscopy of StcE^E447D^ staining of unpermeabilized MCF10A cells cultured on 400 Pa in HPLM or DMEM with or without 15 mM glucose.

**Figure S5:**
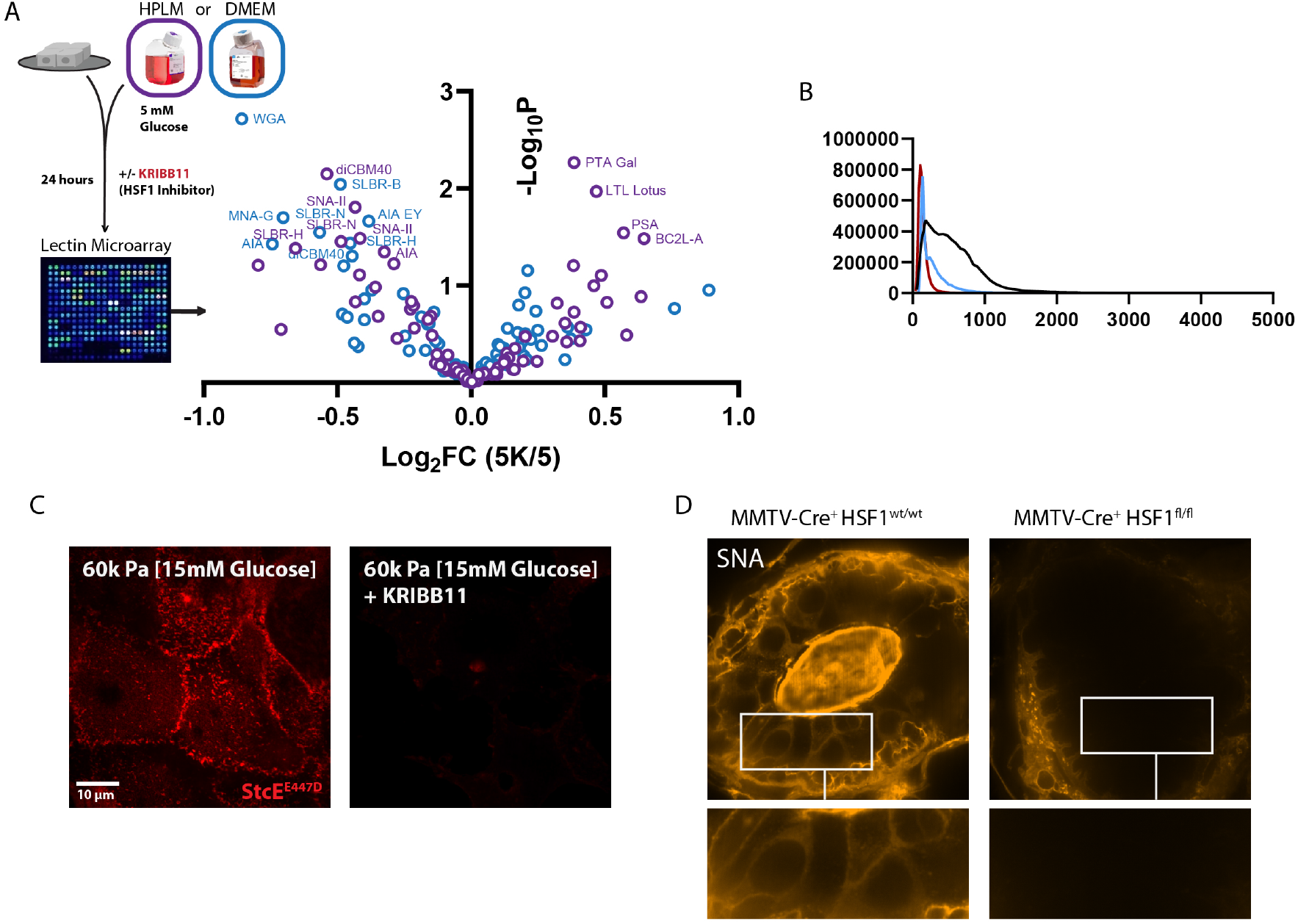
A. Volcano plot depicting relative abundance of glycans from MCF10A cells cultured in HPLM with 5 mM glucose +/- KRIBB11 [2 µM], detected via lectin microarray. (n = 4 biological replicates) B. Associated fluorescent intensity histogram for Figure 5F, KRIBB11 (red), StcE (blue), vehicle (black). C. Representative SoRa confocal microscopy of StcE^E447D^ staining of MCF10A cells on 60kPa in HPLM with 15 mM glucose +/- KRIBB11 [2 uM]. D. SNA staining associated with Figure 5G.

**Figure S6:**
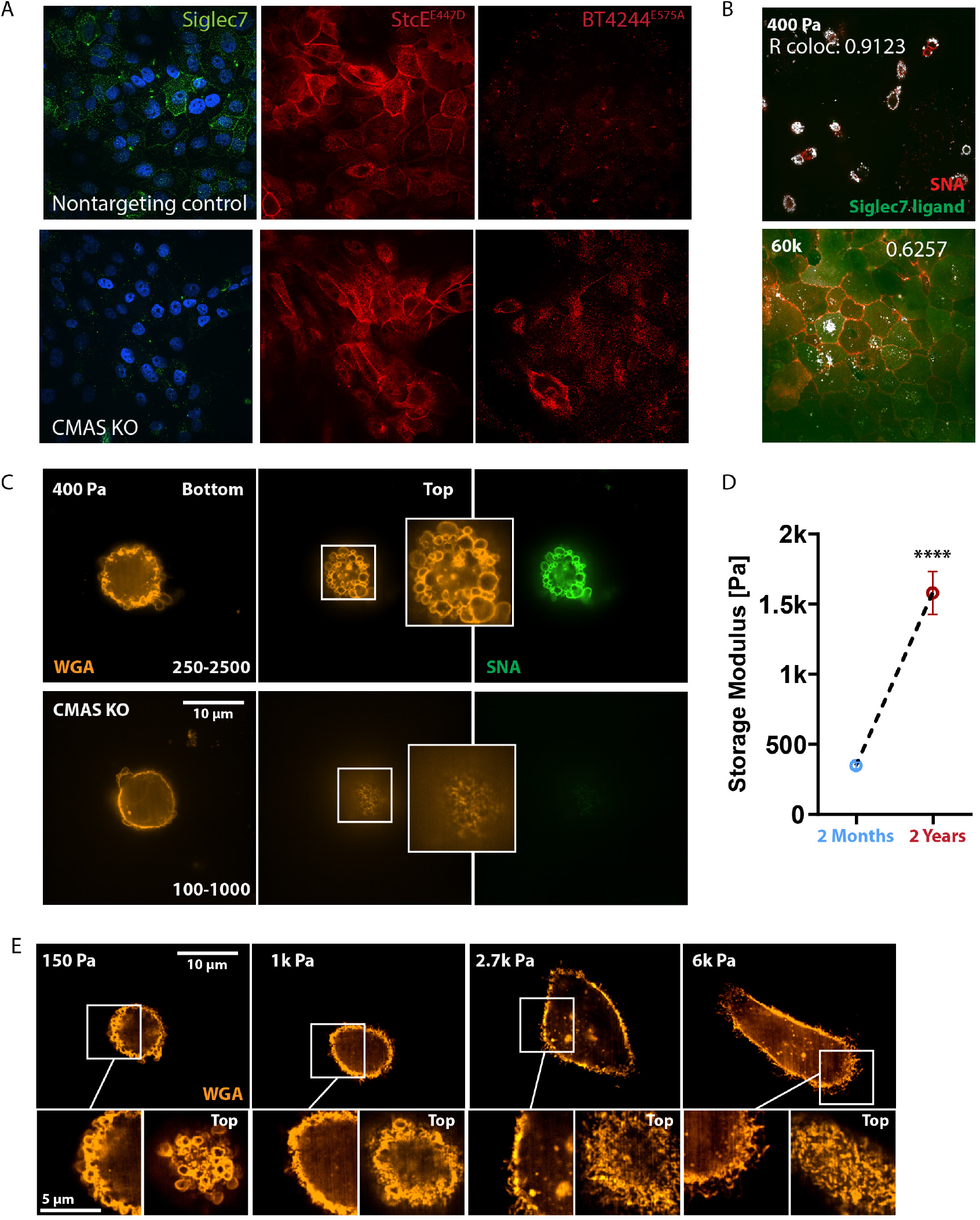
A. Representative confocal microscopy of StcE^E447D^(red), BT4244^E575A^ (Red), Siglec7 Fc (green), and DAPI (blue) staining of non-targeting WT controls and CMAS KO MCF10A cells in HPLM. B. Representative confocal microscopy and colocaization (white) of Siglec7 Fc (green) and SNA (red) staining of MCF10A cells cultured on 400 Pa or 60k Pa in HPLM. C. Representative SoRa confocal microscopy of WGA (orange) and SNA (green) staining of non-targeting WT controls and CMAS KO MCF10A cells on 400 Pa in HPLM. D. Storage modulus of murine mammary glands from 2 month- or 2 year-old female C57BL6/J mice, measured with parallel plate rheology. (n = 5) E. Representative SoRa confocal microscopy of WGA staining of MCF10A cells on 150 Pa, 1k Pa, 2.7k Pa, and 6k Pa in HPLM.

**Figure S7:**
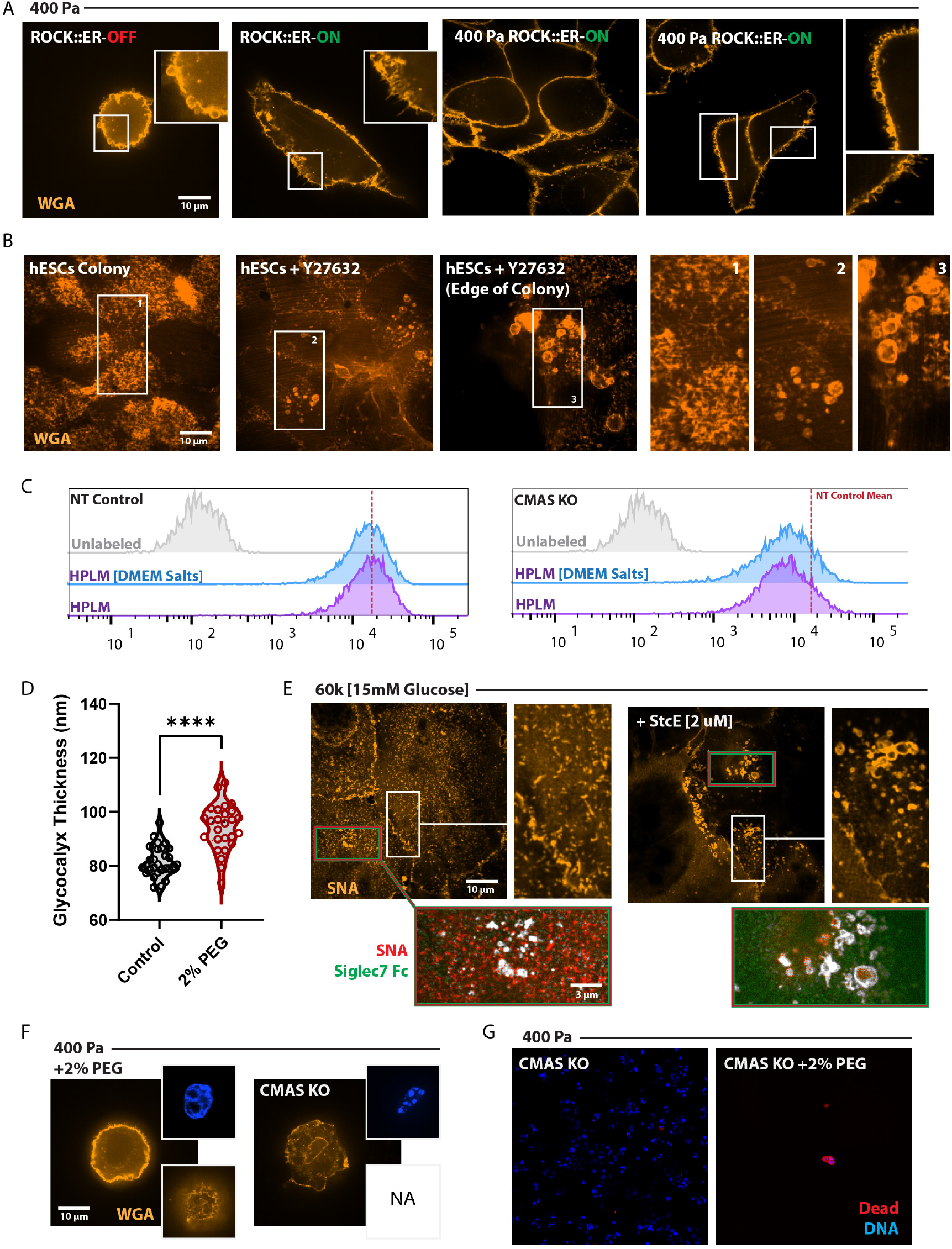
A. Representative confocal microscopy of WGA staining of unpermeabilized ROCK::ER MCF10A cells cultured on 400 Pa with and without 4-HT, single cells and multicellular colonies shown. B. Representative SORA confocal microscopy of WGA staining of hESCs cultured on glass coverslips with and without 10 uM Y27632. C. FACS analysis of WGA staining on ∼ 5k non-targeting WT controls and CMAS KO MCF10A cells on in HPLM or HPLM^DMEM^ ^Salt^. D. SAIM-based quantification of glycocalyx thickness of MCF10A cells in HPLM with and without 2% PEG-400 for 4 hours, results are the mean ± S.D. of at least 13 cells per condition (repeated 2 separate times with similar effects). E. Representative SoRa confocal microscopy and colocaization (white) of Siglec7 Fc (green) and SNA (orang/red) staining of MCF10A cells cultured on 60k Pa +/- StcE [2 uM] for 24 h. (+ StcE condition has fluorescent intensity enhanced 10x) F. Representative SoRa confocal microscopy of WGA (orange) and DAPI (blue) staining of non-targeting WT controls and CMAS KO MCF10A cells on 400 Pa in HPLM with 2% PEG-400 (v/v) for 24 h. G. Representative confocal microscopy of propidium iodide homodimer (red) and calcien-AM (white) staining of non-targeting WT controls and CMAS KO MCF10A cells on 400 Pa in HPLM with 2% PEG-400 (v/v) for 24 h.

## References

1. K. M. Tharp, V. M. Weaver, Modeling Tissue Polarity in Context. J. Mol. Biol. 430, 3613–3628 (2018).

2. M. W. Pickup, J. K. Mouw, V. M. Weaver, The extracellular matrix modulates the hallmarks of cancer. EMBO Rep. 15, 1243–1253 (2014).

3. B. Piersma, M. K. Hayward, V. M. Weaver, Fibrosis and cancer: A strained relationship. Biochim. Biophys. Acta - Rev. Cancer. 1873, 188356 (2020).

4. J. S. Park, C. J. Burckhardt, R. Lazcano, L. M. Solis, T. Isogai, L. Li, C. S. Chen, B. Gao, J. D. Minna, R. Bachoo, R. J. DeBerardinis, G. Danuser, Nature, in press, doi:10.1038/s41586-020-1998-1.

5. V. Papalazarou, T. Zhang, N. R. Paul, A. Juin, M. Cantini, O. D. K. K. Maddocks, M. Salmeron-Sanchez, L. M. Machesky, The creatine–phosphagen system is mechanoresponsive in pancreatic adenocarcinoma and fuels invasion and metastasis. Nat. Metab. 2, 62–80 (2020).

6. T. M. J. Evers, L. J. Holt, S. Alberti, A. Mashaghi, Reciprocal regulation of cellular mechanics and metabolism. Nat. Metab. 2021 34. 3, 456–468 (2021).

7. Y. Wu, M. R. Zanotelli, J. Zhang, C. A. Reinhart-King, Matrix-driven changes in metabolism support cytoskeletal activity to promote cell migration. Biophys. J. 120, 1705–1717 (2021).

8. M. R. Zanotelli, J. Zhang, C. A. Reinhart-King, Mechanoresponsive metabolism in cancer cell migration and metastasis. Cell Metab. 33, 1307–1321 (2021).

9. K. M. Tharp, R. Higuchi-Sanabria, G. A. Timblin, B. Ford, C. Garzon-Coral, C. Schneider, J. M. Muncie, C. Stashko, J. R. Daniele, A. S. Moore, P. A. Frankino, S. Homentcovschi, S. S. Manoli, H. Shao, A. L. Richards, K.-H. Chen, J. ten Hoeve, G. M. Ku, M. Hellerstein, D. K. Nomura, K. Saijo, J. Gestwicki, A. R. Dunn, N. J. Krogan, D. L. Swaney, A. Dillin, V. M. Weaver, Adhesion-mediated mechanosignaling forces mitohormesis. Cell Metab. 33, 1322–1341.e13 (2021).

10. K. M. Tharp, K. Kersten, O. M. Maller, G. A. Timblin, C. Stashko, F. P. Canale, M.-K. Hayward, I. Berestjuk, J. ten Hoeve-Scott, B. Samad, A. J. Ironside, R. Geiger, A. J. Combes, V. M. Weaver, bioRxiv, in press, doi:10.1101/2022.07.14.499764.

11. M. J. Paszek, C. C. Dufort, O. Rossier, R. Bainer, J. K. Mouw, K. Godula, J. E. Hudak, J. N. Lakins, A. C. Wijekoon, L. Cassereau, M. G. Rubashkin, M. J. Magbanua, K. S. Thorn, M. W. Davidson, H. S. Rugo, J. W. Park, D. A. Hammer, G. Giannone, C. R. Bertozzi, V. M. Weaver, The cancer glycocalyx mechanically primes integrin-mediated growth and survival. Nature. 511, 319 (2014).

12. J. C. H. Kuo, J. G. Gandhi, R. N. Zia, M. J. Paszek, Physical biology of the cancer cell glycocalyx. Nat. Phys. 14, 658 (2018).

13. A. Buffone, V. M. Weaver, Don’t sugarcoat it: How glycocalyx composition influences cancer progression. J. Cell Biol. 219 (2020), doi:10.1083/JCB.201910070.

14. G. Wang, S. Kostidis, G. L. Tiemeier, W. M. P. J. Sol, M. R. De Vries, M. Giera, P. Carmeliet, B. M. Van Den Berg, T. J. Rabelink, Shear Stress Regulation of Endothelial Glycocalyx Structure Is Determined by Glucobiosynthesis. Arterioscler. Thromb. Vasc. Biol. 40, 350–364 (2020).

15. M. Y. Pahakis, J. R. Kosky, R. O. Dull, J. M. Tarbell, The role of endothelial glycocalyx components in mechanotransduction of fluid shear stress. Biochem. Biophys. Res. Commun. 355, 228–233 (2007).

16. T. E. Sutherland, D. P. Dyer, J. E. Allen, The extracellular matrix and the immune system: A mutually dependent relationship. Science (80-.). 379 (2023), doi:10.1126/SCIENCE.ABP8964/ASSET/C12C1D83-91B2-42C8-A821-EBA5DBE95EBA/ASSETS/IMAGES/LARGE/SCIENCE.ABP8964-FA.JPG.

17. J. A. A. Gubbels, M. Felder, S. Horibata, J. A. Belisle, A. Kapur, H. Holden, S. Petrie, M. Migneault, C. Rancourt, J. P. Connor, M. S. Patankar, MUC16 provides immune protection by inhibiting synapse formation between NK and ovarian tumor cells. Mol. Cancer. 9, 11 (2010).

18. S. Hong, C. Yu, E. Rodrigues, Y. Shi, H. Chen, P. Wang, D. G. Chapla, T. Gao, R. Zhuang, K. W. Moremen, J. C. Paulson, M. S. Macauley, P. Wu, Modulation of Siglec-7 Signaling Via in Situ-Created High-Affinity cis-Ligands. ACS Cent. Sci. 7, 1338–1346 (2021).

19. J. E. Hudak, S. M. Canham, C. R. Bertozzi, Glycocalyx Engineering Reveals a Siglec-Based Mechanism for NK Cell Immunoevasion. Nat. Chem. Biol. 10, 69 (2014).

20. S. Torrino, E. M. Grasset, S. Audebert, I. Belhadj, C. Lacoux, M. Haynes, S. Pisano, S. Abélanet, F. Brau, S. Y. Chan, B. Mari, W. M. Oldham, A. J. Ewald, T. Bertero, Mechano-induced cell metabolism promotes microtubule glutamylation to force metastasis. Cell Metab. 33, 1342–1357.e10 (2021).

21. M. Chakraborty, K. Chu, A. Shrestha, X. S. Revelo, X. Zhang, M. J. Gold, S. Khan, M. Lee, C. Huang, M. Akbari, F. Barrow, Y. T. Chan, H. Lei, N. K. Kotoulas, J. Jovel, C. Pastrello, M. Kotlyar, C. Goh, E. Michelakis, X. Clemente-Casares, P. S. Ohashi, E. G. Engleman, S. Winer, I. Jurisica, S. Tsai, D. A. Winer, Mechanical Stiffness Controls Dendritic Cell Metabolism and Function. Cell Rep. 34, 108609 (2021).

22. P. Romani, N. Nirchio, M. Arboit, V. Barbieri, A. Tosi, F. Michielin, S. Shibuya, T. Benoist, D. Wu, C. C. T. Hindmarch, M. Giomo, A. Urciuolo, F. Giamogante, A. Roveri, P. Chakravarty, M. Montagner, T. Calì, N. Elvassore, S. L. Archer, P. De Coppi, A. Rosato, G. Martello, S. Dupont, Mitochondrial fission links ECM mechanotransduction to metabolic redox homeostasis and metastatic chemotherapy resistance. Nat. Cell Biol. 2022 242. 24, 168–180 (2022).

23. K. M. Tharp, M. S. Kang, G. A. Timblin, J. Dempersmier, G. E. Dempsey, P.-J. H. Zushin, J. Benavides, C. Choi, C. X. Li, A. K. Jha, S. Kajimura, K. E. Healy, H. S. Sul, K. Saijo, S. Kumar, A. Stahl, Actomyosin-Mediated Tension Orchestrates Uncoupled Respiration in Adipose Tissues. Cell Metab. 27, 602–615.e4 (2018).

24. J. R. Cantor, M. Abu-Remaileh, N. Kanarek, E. Freinkman, X. Gao, A. Louissaint Jr, C. A. Lewis, D. M. Sabatini, Physiologic Medium Rewires Cellular Metabolism and Reveals Uric Acid as an Endogenous Inhibitor of UMP Synthase. Cell. 169, 258–272.e17 (2017).

25. J. R. Cantor, The Rise of Physiologic Media. Trends Cell Biol. 29, 854–861 (2019).

26. N. J. Rossiter, K. S. Huggler, C. H. Adelmann, H. R. Keys, R. W. Soens, D. M. Sabatini, J. R. Cantor, CRISPR screens in physiologic medium reveal conditionally essential genes in human cells. Cell Metab. 33, 1248–1263.e9 (2021).

27. W. Qi, H. A. Keenan, Q. Li, A. Ishikado, A. Kannt, T. Sadowski, M. A. Yorek, I. H. Wu, S. Lockhart, L. J. Coppey, A. Pfenninger, C. W. Liew, G. Qiang, A. M. Burkart, S. Hastings, D. Pober, C. Cahill, M. A. Niewczas, W. J. Israelsen, L. Tinsley, I. E. Stillman, P. S. Amenta, E. P. Feener, M. G. Vander Heiden, R. C. Stanton, G. L. King, Pyruvate kinase M2 activation may protect against the progression of diabetic glomerular pathology and mitochondrial dysfunction. Nat. Med. 23, 753 (2017).

28. N. A. Baird, P. M. Douglas, M. S. Simic, A. R. Grant, J. J. Moresco, S. C. Wolff, J. R. Yates 3rd, G. Manning, A. Dillin, HSF-1-mediated cytoskeletal integrity determines thermotolerance and life span. Science. 346, 360–363 (2014).

29. M. M. Murata, X. Kong, E. Moncada, Y. Chen, H. Imamura, P. Wang, M. W. Berns, K. Yokomori, M. A. Digman, NAD+ consumption by PARP1 in response to DNA damage triggers metabolic shift critical for damaged cell survival. Mol. Biol. Cell. 30, 2584–2597 (2019).

30. J. A. Berg, Y. Zhou, Y. Ouyang, A. A. Cluntun, T. C. Waller, M. E. Conway, S. M. Nowinski, T. Van Ry, I. George, J. E. Cox, B. Wang, J. Rutter, Metaboverse enables automated discovery and visualization of diverse metabolic regulatory patterns. Nat. Cell Biol. 25, 616 (2023).

31. S. Campbell, C. Mesaros, L. Izzo, H. Affronti, M. Noji, B. E. Schaffer, T. Tsang, K. Sun, S. Trefely, S. Kruijning, J. Blenis, I. A. Blair, K. E. Wellen, Glutamine deprivation triggers NAGK-dependent hexosamine salvage. Elife. 10 (2021), doi:10.7554/ELIFE.62644.

32. J. W. Dennis, I. R. Nabi, M. Demetriou, Metabolism, Cell Surface Organization, and Disease. Cell. 139, 1229 (2009).

33. C. Reily, T. J. Stewart, M. B. Renfrow, J. Novak, Glycosylation in health and disease. Nat. Rev. Nephrol. 2019 156. 15, 346–366 (2019).

34. C. Agatemor, M. J. Buettner, R. Ariss, K. Muthiah, C. T. Saeui, K. J. Yarema, Exploiting metabolic glycoengineering to advance healthcare. Nat. Rev. Chem. 2019 310. 3, 605–620 (2019).

35. K. J. Metcalf, M. K. Hayward, E. Berens, A. J. Ironside, C. Stashko, E. S. Hwang, V. M. Weaver, Immunosuppressive glycoproteins associate with breast tumor fibrosis and aggression. Matrix Biol. Plus. 14, 100105 (2022).

36. M. J. Paszek, C. C. Dufort, M. G. Rubashkin, M. W. Davidson, K. S. Thorn, J. T. Liphardt, V. M. Weaver, Scanning Angle Interference Microscopy Reveals Cell Dynamics at the Nano-scale. Nat. Methods. 9, 825 (2012).

37. S. Park, M. J. Colville, C. R. Shurer, L.-T. Huang, J. C.-H. Kuo, J. H. Paek, M. C. Goudge, J. Su, M. P. DeLisa, J. Lammerding, W. R. Zipfel, C. Fischbach, H. L. Reesink, M. J. Paszek, bioRxiv, in press, doi:10.1101/2022.01.28.478211.

38. K. T. Pilobello, P. Agrawal, R. Rouse, L. K. Mahal, Advances in lectin microarray technology: Optimized protocols for piezoelectric print conditions. Curr. Protoc. Chem. Biol. 5, 1 (2013).

39. K. T. Pilobello, D. E. Slawek, L. K. Mahal, A ratiometric lectin microarray approach to analysis of the dynamic mammalian glycome. Proc. Natl. Acad. Sci. U. S. A. 104, 11534 (2007).

40. K. T. Pilobello, L. Krishnamoorthy, D. Slawek, L. K. Mahal, Development of a Lectin Microarray for the Rapid Analysis of Protein Glycopatterns. ChemBioChem. 6, 985–989 (2005).

41. R. Qin, G. Meng, S. Pushalkar, M. A. Carlock, T. M. Ross, C. Vogel, L. K. Mahal, Prevaccination Glycan Markers of Response to an Influenza Vaccine Implicate the Complement Pathway. J. Proteome Res. 21, 1974–1985 (2022).

42. D. Bojar, L. Meche, G. Meng, W. Eng, D. F. Smith, R. D. Cummings, L. K. Mahal, A Useful Guide to Lectin Binding: Machine-Learning Directed Annotation of 57 Unique Lectin Specificities. ACS Chem. Biol. 17, 2993–3012 (2022).

43. C. R. Shurer, J. C. H. Kuo, L. D. M. Roberts, J. G. Gandhi, M. J. Colville, T. A. Enoki, H. Pan, J. Su, J. M. Noble, M. J. Hollander, J. P. O’Donnell, R. Yin, K. Pedram, L. Möckl, L. F. Kourkoutis, W. E. Moerner, C. R. Bertozzi, G. W. Feigenson, H. L. Reesink, M. J. Paszek, Physical Principles of Membrane Shape Regulation by the Glycocalyx. Cell. 177, 1757 (2019).

44. D. J. Shon, S. A. Malaker, K. Pedram, E. Yang, V. Krishnan, O. Dorigo, C. R. Bertozzi, An enzymatic toolkit for selective proteolysis, detection, and visualization of mucin-domain glycoproteins. Proc. Natl. Acad. Sci. 117, 21299–21307 (2020).

45. S. A. Malaker, N. M. Riley, D. J. Shon, K. Pedram, V. Krishnan, O. Dorigo, C. R. Bertozzi, Revealing the human mucinome. Nat. Commun. 2022 131. 13, 1–13 (2022).

46. G. Romain, P. Strati, A. Rezvan, M. Fathi, I. N. Bandey, J. R. T. Adolacion, D. Heeke, I. Liadi, M. L. Marques-Piubelli, L. M. Solis, A. Mahendra, F. Vega, L. J. N. Cooper, H. Singh, M. Mattie, A. Bot, S. S. Neelapu, N. Varadarajan, Multidimensional single-cell analysis identifies a role for CD2-CD58 interactions in clinical antitumor T cell responses. J. Clin. Invest. 132 (2022), doi:10.1172/JCI159402.

47. M. F. Pech, L. E. Fong, J. E. Villalta, L. J. G. Chan, S. Kharbanda, J. J. O’Brien, F. E. McAllister, A. J. Firestone, C. H. Jan, J. Settleman, Systematic identification of cancer cell vulnerabilities to natural killer cell-mediated immune surveillance. Elife. 8, e47362 (2019).

48. Y. Zhang, Q. Liu, S. Yang, Q. Liao, CD58 Immunobiology at a Glance. Front. Immunol. 12, 2212 (2021).

49. M. Nieuwdorp, T. W. Van Haeften, M. C. L. G. Gouverneur, H. L. Mooij, M. H. P. Van Lieshout, M. Levi, J. C. M. Meijers, F. Holleman, J. B. L. Hoekstra, H. Vink, J. J. P. Kastelein, E. S. G. Stroes, Loss of Endothelial Glycocalyx During Acute Hyperglycemia Coincides With Endothelial Dysfunction and Coagulation Activation In Vivo. Diabetes. 55, 480–486 (2006).

50. X. Xie, J. Lu, E. J. Kulbokas, T. R. Golub, V. Mootha, K. Lindblad-Toh, E. S. Lander, M. Kellis, Systematic discovery of regulatory motifs in human promoters and 3′ UTRs by comparison of several mammals. Nature. 434, 338 (2005).

51. M. L. Mendillo, S. Santagata, M. Koeva, G. W. Bell, R. Hu, R. M. Tamimi, E. Fraenkel, T. A. Ince, L. Whitesell, S. Lindquist, HSF1 drives a transcriptional program distinct from heat shock to support highly malignant human cancers. Cell. 150, 549–562 (2012).

52. C. Dai, L. Whitesell, A. B. Rogers, S. Lindquist, Heat shock factor 1 is a powerful multifaceted modifier of carcinogenesis. Cell. 130, 1005–1018 (2007).

53. C. Dai, S. B. Sampson, HSF1: Guardian of Proteostasis in Cancer. Trends Cell Biol. 26, 17–28 (2016).

54. S. V. Himanen, M. C. Puustinen, A. J. Da Silva, A. Vihervaara, L. Sistonen, HSFs drive transcription of distinct genes and enhancers during oxidative stress and heat shock. Nucleic Acids Res. 50, 6102–6115 (2022).

55. A. S. Lee, Glucose-regulated proteins in cancer: molecular mechanisms and therapeutic potential. Nat. Rev. Cancer. 14, 263–276 (2014).

56. S. A. Malaker, K. Pedram, M. J. Ferracane, B. A. Bensing, V. Krishnan, C. Pett, J. Yu, E. C. Woods, J. R. Kramer, U. Westerlind, O. Dorigo, C. R. Bertozzi, The mucin-selective protease StcE enables molecular and functional analysis of human cancer-associated mucins. Proc. Natl. Acad. Sci. 116, 7278–7287 (2019).

57. K. H. Su, J. Cao, Z. Tang, S. Dai, Y. He, S. B. Sampson, I. J. Benjamin, C. Dai, HSF1 critically attunes proteotoxic-stress sensing by mTORC1 to combat stress and promote growth. Nat. Cell Biol. 18, 527 (2016).

58. M. R. Walker, H. L. Goel, D. Mukhopadhyay, P. Chhoy, E. R. Karner, J. L. Clark, H. Liu, R. Li, J. L. Zhu, S. Chen, L. K. Mahal, B. A. Bensing, A. M. Mercurio, O-linked α2,3 sialylation defines stem cell populations in breast cancer. Sci. Adv. 8, eabj9513 (2023).

59. S. Hong, C. Yu, P. Wang, Y. Shi, W. Cao, B. Cheng, D. G. Chapla, Y. Ma, J. Li, E. Rodrigues, Y. Narimatsu, J. R. Yates, X. Chen, H. Clausen, K. W. Moremen, M. S. Macauley, J. C. Paulson, P. Wu, Glycoengineering of NK Cells with Glycan Ligands of CD22 and Selectins for B-Cell Lymphoma Therapy. Angew. Chemie Int. Ed. 60, 3603–3610 (2021).

60. S. J. Moons, G. J. Adema, M. T. G. M. Derks, T. J. Boltje, C. Büll, Sialic acid glycoengineering using N-acetylmannosamine and sialic acid analogs. Glycobiology. 29, 433–445 (2019).

61. S. Neelamegham, L. K. Mahal, Multi-level regulation of cellular glycosylation: From genes to transcript to enzyme to structure. Curr. Opin. Struct. Biol. 40, 145 (2016).

62. S. Chen, P. Agrawal, E. Hernando-Monge, L. Mahal, High-throughput Assay and In Vivo Screen Identify α-2,3-sialylation of CD98 by ST3GAL1 and ST3GAL2 as Essential to Melanoma Survival. FASEB J. 36 (2022), doi:https://doi.org/10.1096/fasebj.2022.36.S1.L7667.

63. M. Yagita, C. L. Huang, H. Umehara, Y. Matsuo, R. Tabata, M. Miyake, Y. Konaka, K. Takatsuki, A novel natural killer cell line (KHYG-1) from a patient with aggressive natural killer cell leukemia carrying a p53 point mutation. Leukemia. 14, 922–930 (2000).

64. H. Xiao, E. C. Woods, P. Vukojicic, C. R. Bertozzi, Precision glycocalyx editing as a strategy for cancer immunotherapy. Proc. Natl. Acad. Sci. 113, 10304–10309 (2016).

65. J. A. Trapani, M. J. Smyth, Functional significance of the perforin/granzyme cell death pathway. Nat. Rev. Immunol. 2, 735–747 (2002).

66. D. H. Raulet, A. Marcus, L. Coscoy, Dysregulated cellular functions and cell stress pathways provide critical cues for activating and targeting natural killer cells to transformed and infected cells. Immunol. Rev. 280, 93–101 (2017).

67. D. R. Croft, M. F. Olson, Conditional regulation of a ROCK-estrogen receptor fusion protein. Methods Enzymol. 406, 541–553 (2006).

68. K. Watanabe, M. Ueno, D. Kamiya, A. Nishiyama, M. Matsumura, T. Wataya, J. B. Takahashi, S. Nishikawa, S. I. Nishikawa, K. Muguruma, Y. Sasai, A ROCK inhibitor permits survival of dissociated human embryonic stem cells. Nat. Biotechnol. 2007 256. 25, 681–686 (2007).

69. M. Guo, A. F. Pegoraro, A. Mao, E. H. Zhou, P. R. Arany, Y. Han, D. T. Burnette, M. H. Jensen, K. E. Kasza, J. R. Moore, F. C. Mackintosh, J. J. Fredberg, D. J. Mooney, J. Lippincott-Schwartz, D. A. Weitz, Cell volume change through water efflux impacts cell stiffness and stem cell fate. Proc. Natl. Acad. Sci. U. S. A. 114, E8618–E8627 (2017).

70. K. Xie, Y. Yang, H. Jiang, Controlling Cellular Volume via Mechanical and Physical Properties of Substrate. Biophys. J. 114, 675–687 (2018).

71. K. M. Flickinger, K. M. Wilson, N. J. Rossiter, A. L. Hunger, T. D. Lee, M. D. Hall, J. R. Cantor, bioRxiv, in press, doi:10.1101/2023.06.04.543621.

72. R. Krishnan, J. A. Park, C. Y. Seow, P. V. S. Lee, A. G. Stewart, Cellular Biomechanics in Drug Screening and Evaluation: Mechanopharmacology. Trends Pharmacol. Sci. 37, 87 (2016).

73. H. Ichise, S. Tsukamoto, T. Hirashima, Y. Konishi, C. Oki, S. Tsukiji, S. Iwano, A. Miyawaki, K. Sumiyama, K. Terai, M. Matsuda, Functional visualization of NK cell-mediated killing of metastatic single tumor cells. Elife. 11, e76269 (2022).

74. O. Y. Wouters, D. T. A. Ploeger, S. M. van Putten, R. A. Bank, 3,4-Dihydroxy-L-Phenylalanine as a Novel Covalent Linker of Extracellular Matrix Proteins to Polyacrylamide Hydrogels with a Tunable Stiffness. Tissue Eng. Part C. Methods. 22, 91–101 (2016).

75. B. A. Bensing, Q. Li, D. Park, C. B. Lebrilla, P. M. Sullam, Streptococcal Siglec-like adhesins recognize different subsets of human plasma glycoproteins: implications for infective endocarditis. Glycobiology. 28, 601–611 (2018).

76. C. Stringari, A. Cinquin, O. Cinquin, M. A. Digman, P. J. Donovan, E. Gratton, Phasor approach to fluorescence lifetime microscopy distinguishes different metabolic states of germ cells in a live tissue. Proc. Natl. Acad. Sci. 108, 13582–13587 (2011).

77. A. E. Y. T. Lefebvre, D. Ma, K. Kessenbrock, D. A. Lawson, M. A. Digman, Automated segmentation and tracking of mitochondria in live-cell time-lapse images. Nat. Methods. 18, 1091–1102 (2021).

78. J. Doellinger, A. Schneider, M. Hoeller, P. Lasch, Sample Preparation by Easy Extraction and Digestion (SPEED) - A Universal, Rapid, and Detergent-free Protocol for Proteomics Based on Acid Extraction. Mol. Cell. Proteomics. 19, 209 (2020).

79. F. Meier, A.-D. Brunner, S. Koch, H. Koch, M. Lubeck, M. Krause, N. Goedecke, J. Decker, T. Kosinski, M. A. Park, N. Bache, O. Hoerning, J. Cox, O. Räther, M. Mann, Online Parallel Accumulation-Serial Fragmentation (PASEF) with a Novel Trapped Ion Mobility Mass Spectrometer. Mol. Cell. Proteomics. 17, 2534–2545 (2018).

80. J. Cox, N. Neuhauser, A. Michalski, R. A. Scheltema, J. V. Olsen, M. Mann, Andromeda: A peptide search engine integrated into the MaxQuant environment. J. Proteome Res. 10, 1794–1805 (2011).

81. J. Cox, M. Mann, MaxQuant enables high peptide identification rates, individualized p.p.b.-range mass accuracies and proteome-wide protein quantification. Nat. Biotechnol. 26, 1367–1372 (2008).

82. J. E. Elias, S. P. Gygi, Target-decoy search strategy for increased confidence in large-scale protein identifications by mass spectrometry. Nat. Methods. 4, 207–214 (2007).

83. M. Choi, C.-Y. Chang, T. Clough, D. Broudy, T. Killeen, B. MacLean, O. Vitek, MSstats: an R package for statistical analysis of quantitative mass spectrometry-based proteomic experiments. Bioinformatics. 30, 2524–2526 (2014).

84. D. J. Puleston, M. D. Buck, R. I. Klein Geltink, R. L. Kyle, G. Caputa, D. O’Sullivan, A. M. Cameron, A. Castoldi, Y. Musa, A. M. Kabat, Y. Zhang, L. J. Flachsmann, C. S. Field, A. E. Patterson, S. Scherer, F. Alfei, F. Baixauli, S. K. Austin, B. Kelly, M. Matsushita, J. D. Curtis, K. M. Grzes, M. Villa, M. Corrado, D. E. Sanin, J. Qiu, N. Pällman, K. Paz, M. E. Maccari, B. R. Blazar, G. Mittler, J. M. Buescher, D. Zehn, S. Rospert, E. J. Pearce, S. Balabanov, E. L. Pearce, Polyamines and eIF5A Hypusination Modulate Mitochondrial Respiration and Macrophage Activation. Cell Metab. 30, 352–363.e8 (2019).

85. M. J. Colville, S. Park, W. R. Zipfel, M. J. Paszek, High-speed device synchronization in optical microscopy with an open-source hardware control platform. Sci. Rep. 9, 12188 (2019).

86. K. T. Pilobello, P. Agrawal, R. Rouse, L. K. Mahal, Advances in Lectin Microarray Technology: Optimized Protocols for Piezoelectric Print Conditions. Curr. Protoc. Chem. Biol. 5, 1–23 (2013).

87. S. Koppolu, L. Wang, A. Mathur, J. A. Nigam, C. S. Dezzutti, C. Isaacs, L. Meyn, K. E. Bunge, B. J. Moncla, S. L. Hillier, L. C. Rohan, L. K. Mahal, Vaginal Product Formulation Alters the Innate Antiviral Activity and Glycome of Cervicovaginal Fluids with Implications for Viral Susceptibility. ACS Infect. Dis. 4, 1613–1622 (2018).

